# Evolution of innate behavioral strategies through competitive population dynamics

**DOI:** 10.1101/2021.06.24.449791

**Authors:** Tong Liang, Braden A. W. Brinkman

## Abstract

Many organism behaviors are innate or instinctual and have been “hard-coded” through evolution. Current approaches to understanding these behaviors model evolution as an optimization problem in which the traits of organisms are assumed to optimize an objective function representing evolutionary fitness. Here, we use a mechanistic birth-death dynamics approach to study the evolution of innate behavioral strategies in a population of organisms *in silico*. In particular, we performed agent-based stochastic simulations and mean-field analyses of organisms exploring random environments and competing with each other to find locations with plentiful resources. We find that when organism density is low, the mean-field model allows us to derive an effective objective function, predicting how the most competitive phenotypes depend on the exploration-exploitation trade-off between the scarcity of high-resource sites and the increase in birth rate those sites offer organisms. However, increasing organism density alters the most competitive behavioral strategies and precludes the existence of a well-defined objective function. Moreover, there exists a range of densities for which the coexistence of many phenotypes persists for evolutionarily long times.

## INTRODUCTION

Foraging for resources—shelter, food, etc.—is one of the fundamental problems an organism must solve in order to survive and reproduce, ensuring the continuity of its species [1]. Many of the strategies organisms employ to solve such problems appear to be instinctual or innate; for example, birds and fish know where to migrate even if they have never visited their migration grounds before [2], and many complex species such as fish, reptiles, birds, and insects have been found to employ search strategies similar to the much simpler *C. Elegans* [3]. The fact that many disparate species have converged upon similar strategies suggests there may be unifying principles that can be captured by simple quantitative models to help explain the emergence of these behavioral patterns.

Perhaps the most well-known candidate principle is “survival of the fittest:” species with adaptations that give them any edge for outcompeting other species in their environments are those that go on to proliferate. On its own this principle is somewhat tautologous, as the organisms deemed fittest for their environment are those observed to survive and proliferate [4]. In order to render this problem well-posed for quantitative modeling, a common approach is to define a “fitness function,” a function of a population’s phenotypic parameters whose output is a proxy for that phenotypic configuration to survive and reproduce in its environment. Often the fitness function is modeled as the fitness of a single representative agent, rather than an entire population. This idea has given rise to modeling evolution as an optimization process that maximizes an organism’s fitness [4], i.e., selects for the phenotype parameters that maximize the fitness function. Fitness functions can range from relatively straightforward and concrete quantities, such as an organism’s rate of producing offspring or the time it takes to find shelter or food [5–8], or they can be relatively abstract, such as the amount of information an organism’s sensory organs convey to its nervous system [9–12]—the idea being that the more efficiently an organism’s nervous system can process information it acquires about its environment, the more successful it will be at finding food, shelter, mates, etc.

Modeling evolution as an optimization process has had some notable successes in predicting features of biological organisms [12–17]; however, some caveats and cautions are in order. Firstly, one can always find an objective function and set of constraints for which an observed solution is optimal [18]. Secondly, there are examples in which evolution is demonstrably not optimal [19], exhibits significant deviations from optimality [16], or can even be repeatedly reversible [20, 21]. These and other issues with modeling evolution as optimization have been discussed extensively by Doebeli *et al.* [4]. Doebeli *et al.* emphasize that the fundamental events in evolution are stochastic birth-death processes, which can give rise to complex evolutionary dynamics that cannot always be captured by optimizing fitness. For instance, an approach that focuses on birth-death events of entire populations can yield the distribution of successful phenotypes, including phenotypes that would be considered “sub-optimal” by optimization approaches yet are able to coexist with phenotypes predicted to be optimal.

In this paper we take the mechanistic approach, simulating the dynamics of a population of organisms with different innate behavioral strategies for exploring their environment in search of locations with favorable conditions for reproduction. In particular, non-consumable resources are distributed randomly throughout the environment, and each organism’s behavioral strategy consists of the set of rates at which the organism will leave locations of each resource level in search of other locations with potentially greater amounts of the resource. Each location in the environment can only support a single individual, resulting in competitive population dynamics. The most competitive phenotypes in our model will be the set of relocation rates that enable organisms to proliferate, allowing these phenotypes to persist in the population over long time scales. Importantly, the birth rates of organisms in our model depend only on an individual’s location within the environment, not that individual’s phenotype (i.e., locations with plentiful resources confer larger birth rates, regardless of the phenotype of the organism). Thus, the reproductive competitiveness of a phenotype is only determined by how that phenotype enables our model organisms to outcompete other phenotypes in discovering high-resource locations. There is no explicit fitness function that the dynamics are assumed or enforced to optimize. However, we show that in the limit of low interaction density we can derive a growth rate that is a function of the hopping rate parameters and serves as an emergent fitness function. However, we also show that in general this function does not describe the most successful phenotypes when interactions between individuals are frequent, or when demographic fluctuations are significant.

This paper is organized as follows: in Results we first define our stochastic model and its formulations as an agent-based simulation and a corresponding analytic description in terms of a stochastic master equation (Mechanistic model of evolutionary dynamics). We then show that the environmental structure and the population dynamics of the simulated model indeed select for particular phenotypes, even though birth rates depend only on spatial location and *not* the phenotype. Importantly, we find that while some components of the phenotype experience selection pressures strong enough to select for a very narrow range of values, others experience relative weak selection pressure, resulting in distributions of nearly equally competitive phenotypes (Distributions of competitive phenotypes are selected by birth-death dynamics). We show that we can understand the origins of weak selection pressures using a minimal mean-field model, in which we can derive the growth rate as a function of phenotype, which acts as an effective fitness function, when organism density is low (A minimal mean-field model explains variability in competitive phenotypes). In this limit we also study how the exploration-exploitation trade-off depends on the scarcity of high-resource sites and their corresponding birth rate (Distribution of resources shapes exploration-exploitation trade-offs). At higher density the interactions between individual conspecifics cannot be neglected and alters the dominant phenotypes: at very high densities there is a reversal in the most competitive phenotypes, and there is a range of intermediate densities at which many phenotypes coexist for evolutionarily long times (Ranges of population density allow for long-lasting phenotype coexistence). We conclude in the Discussion by addressing the interpretation of our results in the context of mechanistic and normative approaches to the evolution of biological structure (Summary and interpretation of results) and comparing our results to those of related studies (Comparisons to related work), as well as discussing the limitations of our approach (Limitations of the study) and future directions for extensions of this work (Future directions).

## RESULTS

### Mechanistic model of evolutionary dynamics

To investigate the evolution of foraging behavioral strategies, we focus on modeling a population of organisms that must navigate their environment in search of locations favorable for reproduction. For simplicity, we will assume the favorability of each location is determined by the amount of a non-depleting “resource” at this location. Additionally, each location has a fixed carrying capacity, allowing only a single individual to occupy the location at any given moment. Organisms are able to detect the resource level only at their current locations, and must decide whether to remain where they are or explore in search of regions with more resources (see Fig. 1A); this is referred to as the “exploration-exploitation trade-off” [22]. In this model the organisms’ movements are not deterministic, but instead each individual will attempt to relocate stochastically at rates determined by the amount of resource at the individual’s current location; the value of these rates is determined by the individual’s phenotype. The resource level at each location is predetermined and sampled from a discrete distribution; see Fig. 1B, depicting a discrete uniform resource distribution. The set of relocation rates (used interchangeably with the term “hopping rates”) for each resource level comprises an organism’s phenotype in this model; see Fig. 1C.

**FIG. 1.**
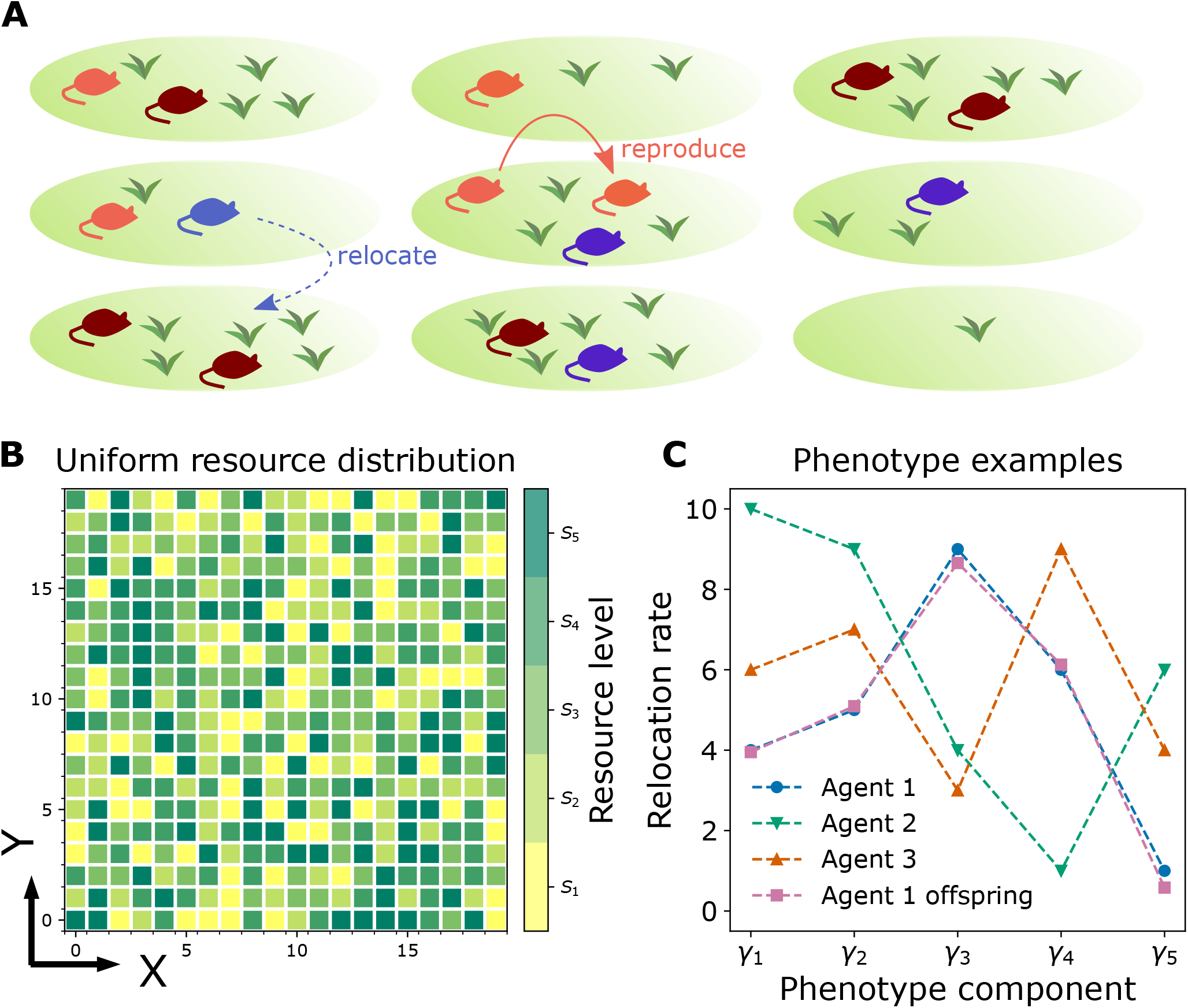
Schematic diagram of the mechanistic evolutionary dynamics model. **A.** The model consists of a population of agents foraging in an environment with a random distribution of resources, in which each agent can either asexually produce an offspring (as shown by the orange arrow) or relocate to one of the vertical or horizontal neighboring locations (as shown by the dashed blue arrow). Different colors here distinguish agents with different phenotypes. **B.** The environment is a two dimensional discrete lattice on which each location has a predetermined resource level *S*_*k*_, *k* ∈ {1, 2, 3, 4, 5}, chosen from a distribution (a discrete uniform distribution shown here). A 20 × 20 subset of the environment is shown here. Each site on the lattice is colored according to its resource level *S*_*k*_. **C.** Each agent has a phenotype vector with 5 components, corresponding to the relocation or hopping rates of the agent at 5 different resource levels. These components are initially chosen randomly for each agent and are shaped over the course of evolution. In the example shown here, agent 2 initially has a relocation rate *γ*_3_ ≈ 4 when it is at a site with resource level *S*_3_. In the agent-based stochastic model, an offspring inherits a noisy copy of its parent’s phenotype, as shown here for the phenotypes of agent 1 and its offspring.

In addition to determining an organism’s relocation rate, the resource level at each site directly determines the organisms’ birth rates at that location. Importantly, the birth rates *do not* depend on the organisms’ phenotype—the only way the phenotype influences the organism’s reproductive success is by improving the organism’s odds of encountering high-resource sites and staying there long enough to produce sufficiently large numbers of offspring. That is, the selective advantage of a phenotype is due only to its dynamical success in the environment, not any intrinsic advantage. For simplicity, we model all reproduction as asexual. In the agent-based model to be defined shortly, the offsprings’ phenotypes are taken to be noisy copies of the parents’ phenotype (see “Agent 1” and “Agent 1 offspring” in Fig. 1C). In the mean-field model discussed in the next section, the offsprings’ phenotypes will be direct copies of the parents’.

The final ingredient the model needs is a means of selection. Typically, one would allow for stochastic death of individuals, such that phenotypes that do not confer a sufficiently competitive advantage will eventually die out, leaving the more competitive phenotypes to dominate the population. However, a technical downside of stochastic death is that extinction becomes not only possible but inevitable, potentially complicating investigation of long-term phenotype distributions. To avoid the technical complications of stochastic death, we instead induce selection by subjecting the population to a periodically-occurring “catastrophe” that randomly culls individuals until the population is reduced to its initial size. In doing so, the population is never at risk for extinction, but selection still occurs because the phenotypes that are able to produce large numbers of offspring have better odds of surviving each catastrophe.

To formalize this schematic picture, we first build an agent-based stochastic model. In this model the population consists of individual organisms, or “agents,” each of which is assigned a location within a 2*d* environment, modeled as a discrete lattice, and the values of its hopping rates *γ* at each resource level. In the simulations we investigate here, we assume the resource level of site at location x, labeled *s*_x_, is drawn from 5 distinct resource levels, *s*_x_ ∈ {*S*_1_, *S*_2_, *S*_3_, *S*_4_, *S*_5_}. Thus each agent has 5 corresponding hopping rates *γ* = {*γ*_1_, *γ*_2_, *γ*_3_, *γ*_4_, *γ*_5_}, one for each of the five resource levels. The values of each of these hopping rates are initially chosen randomly from the range [*γ*_min_, *γ*_max_], where *γ*_min_ is the minimum rate at which an organism relocates and *γ*_max_ is the fastest rate an organism can relocate—we interpret this upper bound on relocation rates as being set by metabolic constraints that we do not model or allow to change during the evolution of the population.

The state of the population of agents is updated in continuous time. At any given moment, an agent can either relocate to or asexually give birth to an offspring into one of the vertical or horizontal neighboring locations, as schematically shown in Fig. 1A (because only one agent is permitted per site, an agent cannot give birth to a child on its current site). The per-capita birth rate of an agent, *b*(*s*_x_), is a function of the local resource level *s*_x_ at location x. Because the resources are not depleted in this model the actual numerical values of the *s*_x_ are unimportant; the meaningful values are the birth rates *b*(*s*_x_). However, we assume that the values of the birth rates are ordered according to resource level, such that a site x with resource level *s*_x_ = *S*_1_ has the lowest birth rate and a site with resource level *s*_x_ = *S*_5_ has the highest birth rate. Again, we stress that the birth rates are entirely independent of an agent’s phenotype *γ*.

Agents with very different phenotypes will navigate the environment in different ways. An agent with a small value of *γ*_1_, for example, will tend to remain in sites of resource level *S*_1_ for long periods of time, while agents with large values of *γ*_1_ will tend to leave sites of this resource level rather quickly. Because the birth rate in sites of resource level *S*_1_ is low, we might expect that agents with small *γ*_1_ may be less successful at proliferating than agents with larger *γ*_1_, and will therefore be less likely to survive the periodic catastrophes that occur, creating a selection pressure toward larger values of *γ*_1_. Similarly, agents with small (large) values of *γ*_5_ will tend to remain in these high-resource sites for long (short) periods of time. Agents with small *γ*_5_ may therefore produce more offspring because *b*(*S*_5_) is the highest, increasing their odds of surviving the catastrophes. This line of thinking makes it seem intuitively “obvious” that the most competitive phenotype ought to have large hopping rates out of low resource sites and minimal hopping out of the highest resource sites. Moreover, we might expect we can identify an objective function that yields this phenotype as its optimum. In the next sections we present the results of our stochastic simulations as well as the numerical solutions of the mean-field model, showing that this intuition is largely correct when the population density is low—in which case we can indeed derive an effective objective function—and when the distribution of resource levels is uniform, but this picture is not always correct when high resource sites are rare or population density is high.

### Distributions of competitive phenotypes are selected by birth-death dynamics

For the results presented in this section, we simulate our agent-based model using the Gillespie algorithm [23], starting with 500 agents on a 128 × 128 2*d* lattice. This corresponds to an initial population density of 3.1%. Each lattice site belongs to one of the 5 resource levels {*S*_1_, *S*_2_, *S*_3_, *S*_4_, *S*_5_}, which is drawn from a discrete uniform distribution. Initially, each agent is assigned a spatial location x = (*x, y*) and a phenotype vector *γ* = (*γ*_1_, *γ*_2_, *γ*_3_, *γ*_4_, *γ*_5_) uniformly at random. Catastrophes occur after a simulation runtime *T*, which defines an iteration of the simulation. While the catastrophe randomly picks which agents to cull, it always reduces the total population back to 500 agents. In Distribution of resources shapes exploration-exploitation trade-offs we consider non-uniform distributions, and in Ranges of population density allow for long-lasting phenotype coexistence we investigate different lattice sizes as a means of changing the initial population density. The details of the simulation setups are presented in Methods Section.

FIG. 2A-D displays the evolution of the distribution of phenotypes over several thousand catastrophes, with snapshots at (A) 0, (B) 50, (C) 500, and (D) 20000 iterations; these distributions represent averages over 20 trials of the dynamics starting from the random initial conditions generated from the same distribution. The initially uniform distributions of each of the five hopping rates (phenotype components) are gradually reshaped by births and deaths. At long times, the distributions are approximately stable; we do not expect the distributions to converge to atomic distributions concentrated at a single value for each *γ*_*k*_ due to the reproduction noise that accompanies each birth. In Fig. 2E we plot the evolution of the mean value of each hopping rate over the 20000 catastrophes, averaged over 20 trials. The shaded regions of the plot correspond to the width of the distributions (the standard deviation of each hopping rate component).

**FIG. 2.**
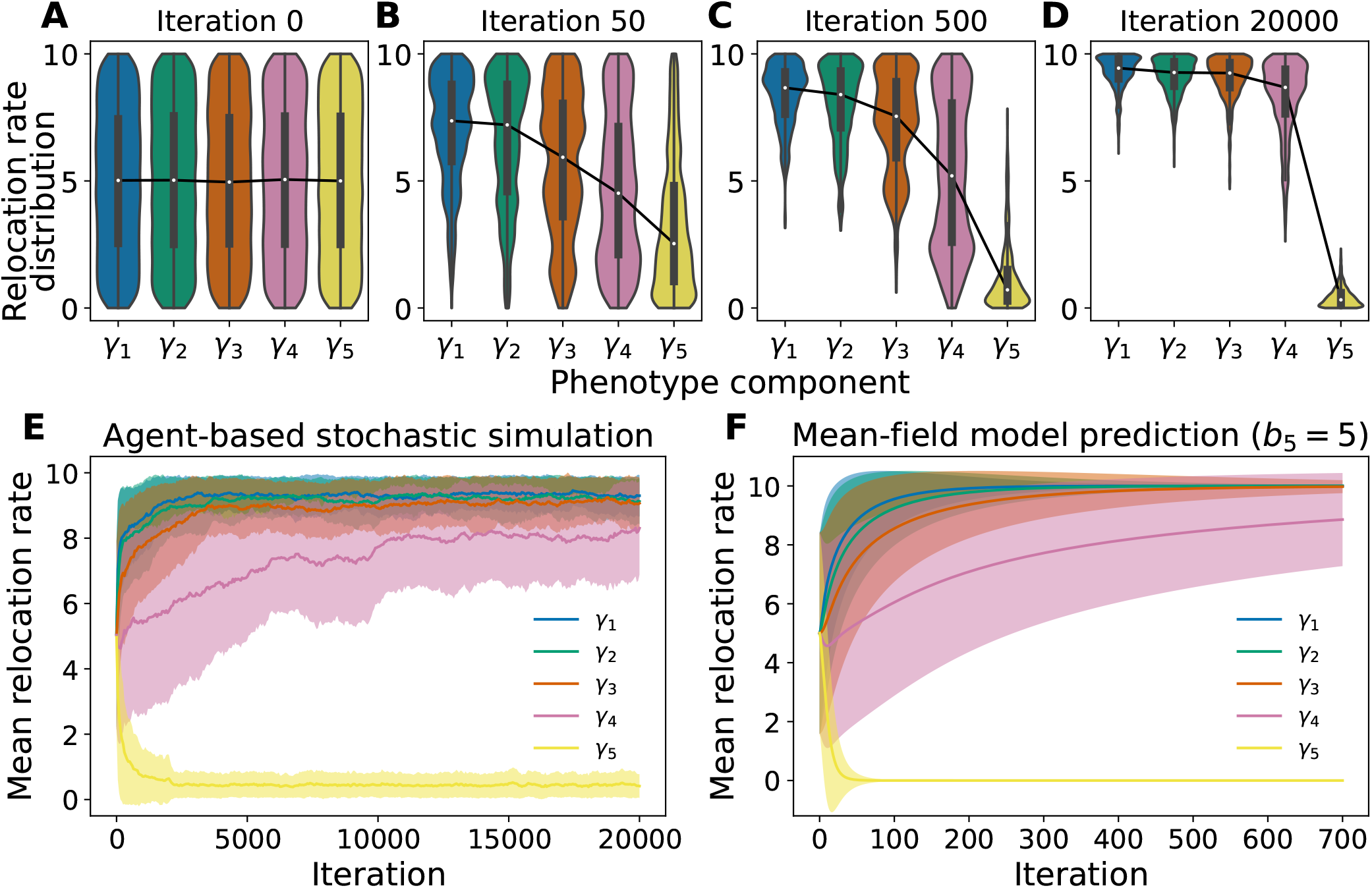
Emergent competitive phenotypes from evolutionary birth-death dynamics. **A**-**D**. Violin plots of distributions of phenotype components in the agent-based stochastic simulations at iterations 0, 50, 500, and 20000, respectively. Each violin plot combines data from 20 independent trials. The connected white dots are the median values of each relocation rate component, and the black boxes confine the first and third quartiles of each phenotype component (i.e., box plot). The colored areas are the kernel density estimations of the underlying distributions. Panel A shows an uniform relocation rate distribution for each phenotype component as the initial condition at iteration 0. The phenotype distributions evolve as the birth-death dynamics happens in the agent-based model and approach a steady shape after around 20000 iterations as shown in panel D. **E** The evolutionary trajectory of each phenotype component, averaged over 20 independent trials. The shaded area represents the standard deviations of phenotype components. The stochastic birth-death dynamics selects out competitive phenotypes whose first three phenotype components *γ*_1,2,3_ saturate at hopping rate of around 9, close to the maximum possible hopping rate of *γ*_max_ = 10, and whose last phenotype component *γ*_5_ saturates at hopping rate around 0.5, close to the minimum possible hopping rate of *γ*_min_ = 0. In contrast, the fourth component *γ*_4_ approaches a value around 8 with a relatively large variability throughout the simulation. **F.** Predictions of the mean relocation rate of each phenotype component and their standard deviations using the coupled 5-state mean-field model, discussed in A minimal mean-field model explains variability in competitive phenotypes. Note that the mean-field model uses a slightly different birth rate *b*_5_ = 5, while the stochastic simulations use *b*_5_ = 4, and the 5-state mean-field model converges at a much shorter timescale (700 iterations vs. 20000 iterations). These quantitative differences between the full agent-based model and the simplified 5-state model are discussed in detail in the Discussion and Supplementary information.

We see that the long-time distribution appears to favor high hopping rates in sites with resource levels *S*_1_ to *S*_3_ and low hopping rates for sites with the highest resource level *S*_5_, reflecting our intuition that the most competitive phenotypes would be those that result in agents leaving their location as fast as possible when at low resource sites and staying put as long as possible when at high resource sites. Notably, however, the distribution of the hopping rates *γ*_4_ at the second-highest resource level *S*_4_ is not narrow, but spans much of the entire possible range of hopping rate values. This indicates that the selection pressure on this phenotype component is relatively weak compared to the selection pressure on the other components. While the average value of *γ*_4_ is closer to *γ*_max_ than *γ*_min_, the large variance demonstrates that the most competitive phenotype does not conform tightly to our intuitive predictions, and the situation is more complex than might be naively expected. To gain insight into our results and investigate how selection pressure depends on our environments’ properties, namely the distribution of resources and the values of the birth rates tied to each resource level, we develop a deterministic minimal mean-field model that enables us to derive an effective objective function in a low-density limit. This fitness function predicts the optimal phenotypes and explains the lack of selection pressure on *γ*_4_ observed in our stochastic simulations.

### A minimal mean-field model explains variability in competitive phenotypes

Agent-based models are often difficult to formulate directly as mathematical models, owing to the combination of having individual with private properties (here, their phenotype) and the fact that agents that may be born or culled at any time. However, a formalism that can approximate the agent-based model outlined here is a master equation formalism that keeps track of the probability of configurations of the number of agents with each phenotype at each spatial location. From such a formalism one can then derive a system of differential equations for the expected number of agents with each phenotype and location. Such a model serves as the starting point for the reduced model we investigate in this section. We focus only on the reduced mean-field model here; see the Methods for details on the full master equation model and the approximate reduction to the minimal mean-field model described below.

When agent density is low, such as in the simulations of the previous section (initial density of 3.1%), we may assume that events in which an agent is blocked from relocating to another site by another occupying agent are rare and may be neglected. In this approximation the growth dynamics of the different phenotypes decouple, and we can describe the dynamics of the population as a set of linear differential equations describing the expected number of agents at each site. To simplify the mean-field model further, we use the fact that each site belongs to one of the five resource levels and only count the number of agents at each level, eliminating the spatial dependence of the environment. This reduces the model to a 5-state model whose solution is the time course of the expected number of agents of each phenotype at each resource level; see the Methods for a detailed derivation of this reduction. The resulting system of differential equations reads

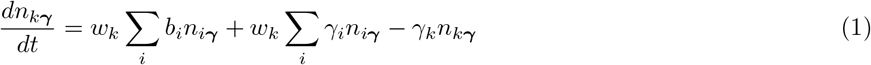

where *n*_*kγ*_ is the expected number of agents at a site with resource level *k* and phenotype *γ*. The parameter *b*_*k*_ is the per-capita birth rate of resource level *k*, a constant of the environment, and *γ*_*k*_ is the hopping rate out of state *k*, determined by the phenotype. Again, we note that in this approximation the dynamics of different phenotypes are decoupled, such that there is a system of differential equations of the form Eq. (1) for each phenotype *γ*. For practical calculations we must discretize the phenotype space, giving a finite system of differential equations. Finally, the weights *w*_*k*_ correspond to the probability that a site in the agent-based model has resource level *S*_*k*_. For an uniform resource distribution, which we investigate first, *w*_*k*_ = 1/5 for all *k* = 1 to 5.

The first term, 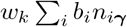, in Eq. (1) is the number of new agents being birthed into state *k* by agents in all other states. The second term 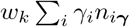 is the number of agents hopping into resource level *k* from all other resource levels, formally including resource level *k* itself to account for the possibility of an agent hopping from a site of resource level *k* to a neighboring site that also has resource level *k*. The last term −*γ*_*k*_*n*_*kγ*_ is the number of agents hopping out of resource level *k*.

The system of differential equations (1) can be written down in matrix form as 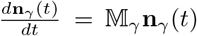, which has a solution in terms of the matrix exponential, 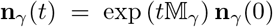. The between-catastrophe dynamics of each phenotype are therefore dominated by the eigenvectors and eigenvalues of 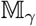. Although the dynamics of the phenotypes are independent, they are coupled through the periodic catastrophes, which reduce the population down to its original size after a time *T*. Let Λ_*γ*_ be the maximum eigenvalue of 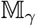, which we interpret as a function of the phenotype parameters *γ* = (*γ*_1_, *γ*_2_, *γ*_3_, *γ*_4_, *γ*_5_). In the Methods we show that the matrix 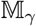 has purely real eigenvalues and only the maximum eigenvalue Λ_*γ*_ is positive. Therefore, the phenotype *γ* that yields with the largest maximum eigenvalue Λ_*γ*_ overall will be the only phenotype to survive after a large number of catastrophes, with coexistence possible only if different phenotypes have the same value of Λ_*γ*_. Because the hopping rates *γ* are just parameters in this formulation of the model, the eigenvalues of 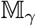 will simply be functions of the phenotype parameters (as well as environmental parameters), and therefore the maximal eigenvalue serves as an *effective objective function* of the system in the low-density limit.

If the maximal eigenvalue of this minimal model acts like an effective objective function, then we would expect the lack of selection pressure on the hopping rate *γ*_4_ to be reflected in an insensitivity of this eigenvalue to varying *γ*_4_, i.e., there would be a range of values of *γ*_4_ for which the maximum eigenvalue approximately takes on its largest value. This is indeed what we find *but only when the birth rate in the highest resource site b*_5_ *is close to a particular critical value B*_5*c*_. As shown in Supplementary Fig. 1, when *b*_5_ > *B*_5*c*_ we find that *γ*_4_ → *γ*_max_ with little sensitivity in the eigenvalue. When *b*_5_ < *B*_5*c*_ and is close to *b*_4_, the birth rate in sites of resource level *S*_4_, we instead find that *γ*_4_ → *γ*_min_, again with little variation in the maximal eigenvalue. This makes sense: if the birth rates *b*_4_ and *b*_5_ are close then there is little advantage in leaving a site with resource level *S*_4_ to search for a site of level *S*_5_. That is, the maximal eigenvalue predicts a transition from *γ*_4_ = *γ*_min_ to *γ*_4_ = *γ*_max_ as *b*_5_ increases from *b*_4_ and crosses *B*_5*c*_. Near the transition at *b*_5_ ≈ *B*_5*c*_ the maximum of the eigenvalue is less sensitive to *γ*_4_—equivalently, fluctuations in *γ*_4_ are large, as is often observed near phase transitions and bifurcations [24], such that there is coexistence of a range of *γ*_4_ values near this transition, at least in a finite population.

Fig. 3A, upper panel, demonstrates that the predicted transition occurs in both our minimal mean-field model and our full agent-based simulations. The critical value *B*_5*c*_ is different for the full stochastic simulation and the mean-field model, but this is to be expected due to the simplifications of the mean-field model as well as the lack of demographic noise. As seen in Fig. 3A, upper panel, the distributions of *γ*_4_ are indeed narrower when *b*_5_ is far from the critical value *B*_5*c*_, reflecting that the selection pressure indeed increases when there is a clear advantage to exploring (*b*_5_ > *B*_5*c*_) versus exploiting (*b*_5_ < *B*_5*c*_). In principle, we might expect to see similar transitions in the other hopping rates *γ*_1_, *γ*_2_, and *γ*_3_ if we adjust the value of *b*_4_, *b*_3_, etc. Instead of varying these parameters to investigate whether such transitions exist, we shift focus to another ecologically important scenario in which these transitions will occur: the effect of the distribution of resources throughout the environment.

**FIG. 3.**
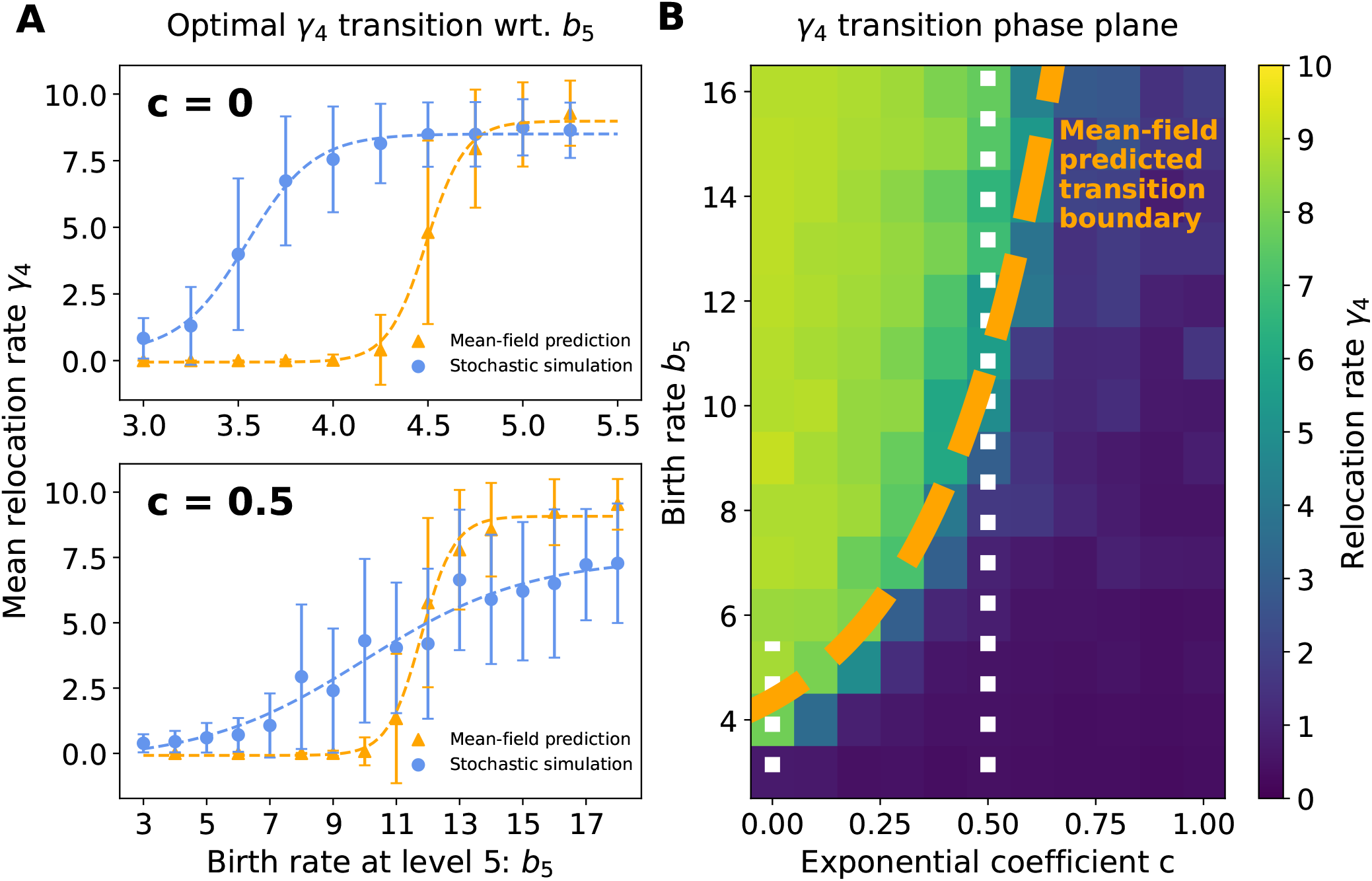
The most competitive phenotypes are shaped by the rarity of the highest-resource sites. The abundance of the five resource levels distributed with frequency *w*_*k*_ ∝ *e*^−*ck*^, where *k* ∈ {1, 2, 3, 4, 5} and the exponential coefficient *c* determines the rarity of levels. The most competitive phenotype changes as a function of birth rates and resource rarity: **A, top.** For a uniform distribution of resources (*c* = 0), the fourth component of the most competitive phenotype *γ*_4_ transitions from low to high as the birth rate *b*_5_ increases. All other phenotype components remain steady as *γ*_1,2,3_ → *γ*_max_ and *γ*_5_ → *γ*_min_. The agent-based stochastic simulation and the mean-field prediction transitions are qualitatively similar, though the transition occurs at different values of *b*_5_. **A, bottom.** A similar transition in *γ*_4_ occurs for a resource distribution with exponential coefficient *c* = 0.5, which had rare high resource level sites. The transition persists but at a higher birth rate *b*_5_. **B.** A heatmap plot of the fourth phenotype component *γ*_4_ as a function of the exponential coefficient *c* and the birth rate *b*_5_. The orange dashed line is the predicted transition boundary from the mean-field model, where *γ*_4_ = 5 maximizes the effective fitness function derived in the low-density limit. The two white dotted lines show the range of *b*_5_ for *c* = 0 and 0.5 in panel A.

### Distribution of resources shapes exploration-exploitation trade-offs

We explore the effect of resource scarcity on the most competitive phenotypes in our model by varying the frequency of the resource levels within the random environment. We do so by randomly assigning the resource level *S*_*k*_ to location *x* with a probability *w*_*k*_ = *Ae*^−*ck*^, where 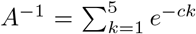 is the normalization constant and *c* is a parameter that determines the steepness of the distribution. The factor *w*_*k*_ is the same factor that appears in Eq. (1). Note that *c* = 0 results in *w*_*k*_ = 1/5, which is the original uniform distribution considered in the previous sections; a positive *c* results in a distribution that resource level rarity increases with *k*, that is, the high resource levels become rarer than the low resource levels.

As shown in Fig. 3A, lower panel, our agent-based simulations predict that the transition from an average value of 〈*γ*_4_〉 = *γ*_min_ = 0 to *γ*_max_ = 10 still exists when *c* is, e.g., 0.5, but occurs at a much larger value of *b*_5_ ≈ 11. Indeed, as seen in Fig. 3B, there exists a transition line in the *b*_5_-*c* plane that separates 〈*γ*_4_〉 ≈ *γ*_min_ = 0 from *γ*_max_ = 10. The orange dashed curve is the boundary predicted by evaluating the maximal eigenvalue of the low-density model Λ_*γ*_ as *c* varies from 0 to 1 and solving for the value of *b*_5_ that yields *γ*_4_ = (*γ*_max_ + *γ*_min_)/2 = 5 as the most competitive phenotype. This curve represents the line along which the population achieves a balance of the exploration-exploitation costs.

Intuitively, agents staying at low resource levels need sufficient incentives to leave their current locations and start to explore new sites, at the risk of missing out reproductive opportunities while exploring. These results demonstrate that rarity of high resource sites can drive selection of phenotypes that settle for sites with lower resource levels.

Importantly, here the rarity of sites is only due to the intrinsic environmental distribution of resources. However, high resource sites could also become *dynamically* rare and unavailable as they become occupied, preventing multiple individuals from exploiting the same sites. So far we have been working within a low-density regime, such that it is relatively rare for organisms’ movements to be blocked by other individuals, and such that a sufficient fraction of high resource sites remain available for discovery and occupation. We now turn to investigating how the competitive phenotypes change with organism density.

### Ranges of population density allow for long-lasting phenotype coexistence

In order to investigate how population density impacts the most competitive phenotype changes, we return to our agent-based stochastic simulations with a uniform resource distribution. The simulations presented previously had 500 agents on a 128 × 128 2*d* lattice, resulting in an initial occupancy level of 3.1%. To study the effect of population density we keep the initial number of agents fixed and decrease the lattice size *L* × *L* to *L* = 96, 80, 64, 48, 40, 35, 32, 28, 26, 25, 24, and 23, giving initial occupancies of 5.4%, 7.8%, 12.2%, 21.7%, 31.2%, 40.8%, 48.8%, 63.8%, 74.0%, 80.0%, 86.8%, and 94.5%, respectively. One could of course also vary density by increasing the initial agent number and keeping the lattice size at 128 × 128, but increasing the number of agents dramatically increases the simulation load.

For each density we ran 20 independent trials of the agent-based stochastic simulation, the results of which are summarized in Fig. 4. We found that as density—and hence interactions among agents—increases, the most competitive phenotypes also change. Notably, these changes qualitatively mimic the effects of increasing the rarity of high resource sites: the mean hopping rate 〈*γ*_4_〉 saturates at a value *γ*_min_ < 〈*γ*_4_〉 ≈ 5 < *γ*_max_ with a large variance at 12.2% initial density, and drops to 〈*γ*_4_〉 ≈ *γ*_min_ with small variance at higher densities.

**FIG. 4.**
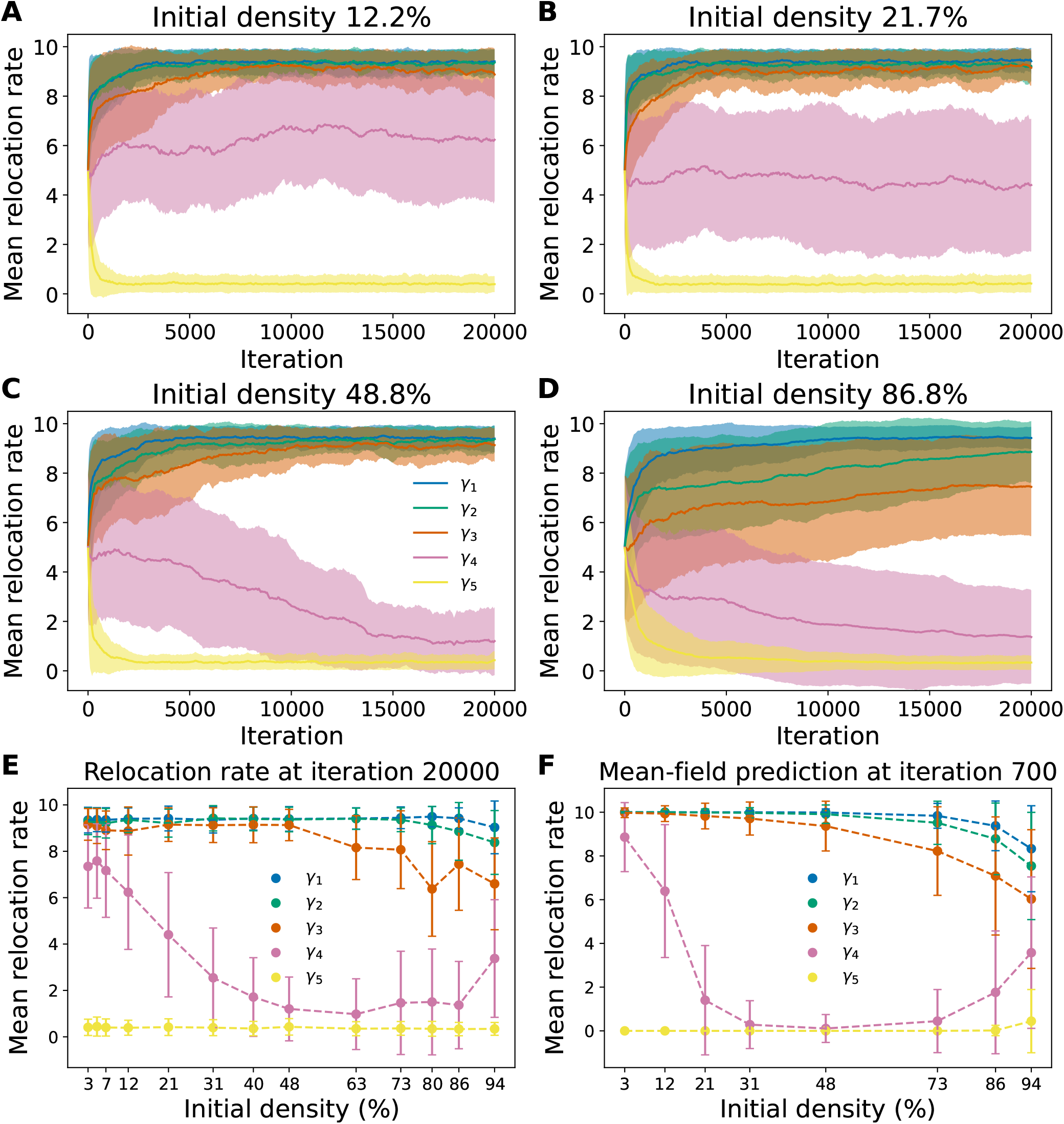
Higher initial densities alter the most competitive phenotypes. **A-D.** The evolution course for populations of agents starting with different initial densities: 12.2%, 21.7%, 48.8%, and 86.8%, respectively; c.f. Fig. 2E at 3.1% density. The initial density is varied by fixing the initial population size at 500 and decreasing the environment size *L* × *L* with *L* = 128, *L* = 64, 48, 32, and 24. We observe a systematic shift of the most competitive phenotypes; in particular, the fourth phenotype component *γ*_4_ decreases as the initial density increases. **E.** Mean values of *γ*_4_ after 20000 iterations in the agent-based model as a function of the initial density, showing the change in the most competitive phenotype. At ultrahigh densities, e.g. 94%, there is no space for agents to move or give birth, resulting in little change in the initially uniform phenotype distribution. **F.** Mean-field model predictions of the most competitive phenotypes as a function of initial density after 700 iterations. A birth rate of *b*_5_ = 5 is used in the mean-field model so that 〈*γ*_4_(*t*)〉 converges to 10 at low densities.

The results observed in our stochastic simulations are replicated by a 5-state mean-field model that incorporates the effects of site occupation:

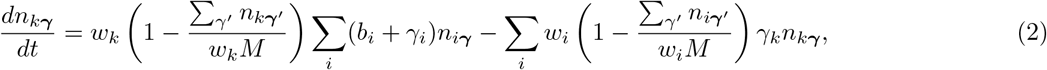

where *n*_*kγ*_ is again the number of agents at sites of resource level *k* and *M* = *L*^2^ is the total number of sites, with *w*_*k*_*M* the expected number of sites of resource level *k*, which sets a maximum occupancy for each state. Importantly, this model is nonlinear, and unlike the low-density approximation, all phenotypes are dynamically coupled. Not only does this preclude writing down an analytical solution, but the dynamical coupling between phenotypes also suggests there may be no effective objective function of a single arbitrary set of the phenotype parameters *γ* = (*γ*_1_, ..., *γ*_5_) that predicts the most competitive phenotype simply by maximizing it with respect to these values.

As in the low-density model, we solve Eq. (2) for a discretized set of phenotypes. Solving the high density model is comparatively computationally demanding due to the necessity of solving for all phenotypes simultaneously, but is feasible for the set of phenotypes chosen here. As shown in Fig. 4F, the mean-field predictions are consistent with our full stochastic simulations, and we indeed observe changes in the most competitive phenotype as a function of initial density *N*_0_/*M*. At low densities the value of 〈*γ*_4_〉 are close to *γ*_max_ (after 700 catastrophes), and as *N*_0_/*M* increases we see a coexistence region in which multiple phenotypes are present at *T*_obs_, before 〈*γ*_4_〉 drops to *γ*_min_ at densities *N*_0_/*M* ≿ 20%. As *N*_0_/*M* increases towards 1, we observe an increase in 〈*γ*_4_〉. This extreme high-density limit represents the scenario in which the population is so dense that there is little room for agents to give birth or move, leading to an extremely slow change in the composition of the phenotypes. In the extreme limit *N*_0_/*M* = 1 we have 〈*γ*_4_(*t*)〉 = (*γ*_max_ + *γ*_min_)/2 for all times *t*, due to the facts that phenotypes are initially present in equal proportions and there is no room for any agent to move or give birth.

The mean-field model is useful for dissecting the dynamics of the phenotype changes. Initially, we hypothesized that the dominance of *γ*_4_ = *γ*_min_ phenotypes at high densities is due to the *S*_5_ resource sites filling up, leaving any *γ*_4_ = *γ*_max_ agents that did not find an *S*_5_ site to search fruitlessly for resource-level 5 sites instead of producing offspring, allowing the *γ*_4_ = *γ*_min_ phenotypes to gain the reproductive advantage. However, as shown in Fig. 5, it turns out that early in the evolutionary dynamics the *γ*_4_ = *γ*_max_ agents lag behind the other phenotypes, as the *γ*_4_ = *γ*_max_ agents leave *S*_4_ sites more frequently and explore the neighboring locations while agents with lower *γ*_4_ gain an advantage by tending to stay at *S*_4_ longer and produce offspring at a relatively high birth rate *b*_4_. At low densities the *γ*_4_ = *γ*_max_ agents eventually discover *S*_5_ sites and their reproduction rate is able to overcome their slow reproductive start. However, at higher densities the *γ*_4_ = *γ*_max_ agents are increasingly less able to overcome the other phenotypes’ head start, and are eventually outcompeted completely by the *γ*_4_ = *γ*_min_ phenotypes at sufficiently high densities.

**FIG. 5.**
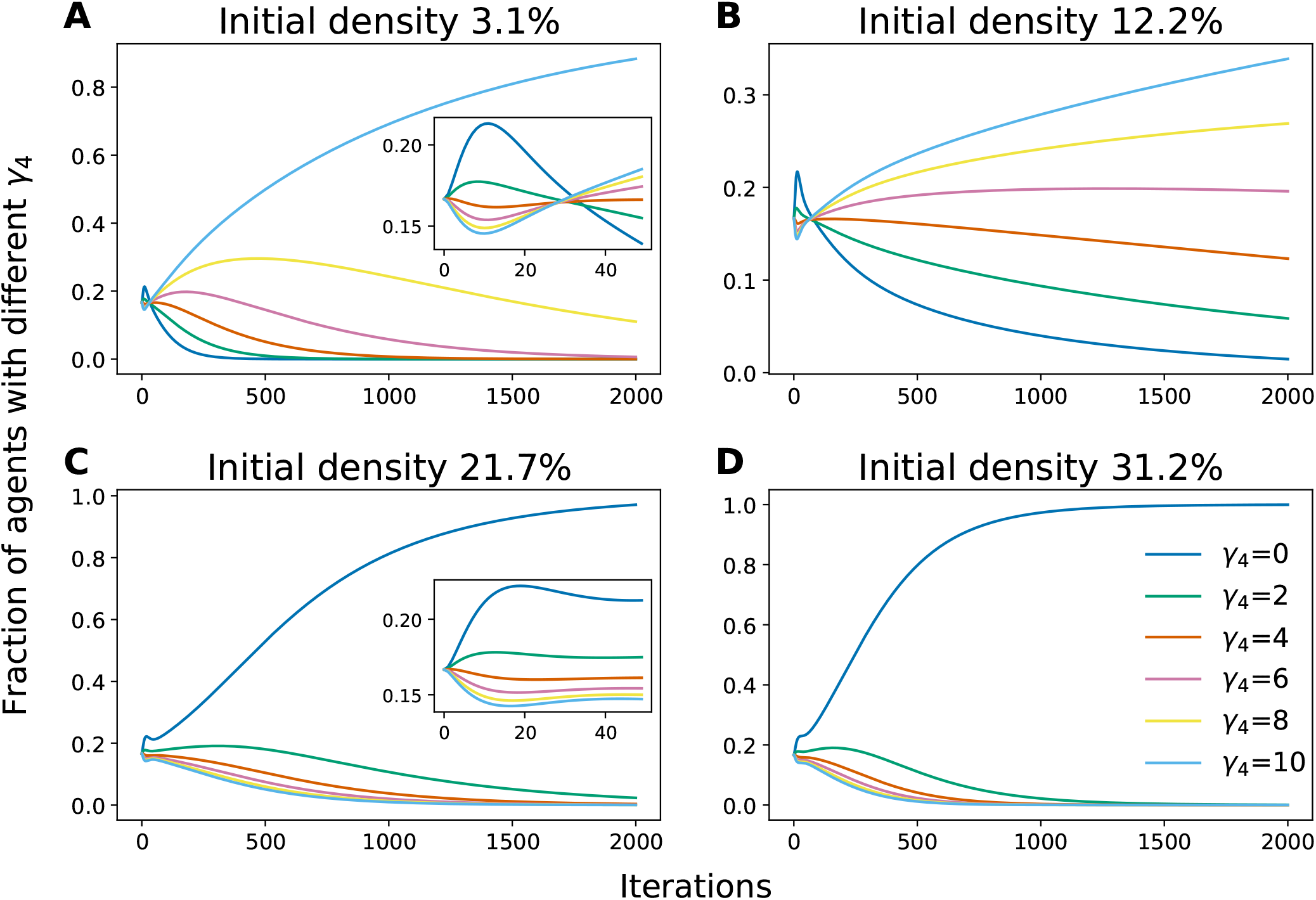
Numerical solutions of the coupled mean-field model at different initial densities. **A-D** show the evolution of the fraction of the population with each value of *γ*_4_ in the mean-field model; insets at 3.1% and 21.7% initial densities show a zooming in of the initial transient dynamics from iteration 0 to 50. Initially in the evolution, *γ*_4_ = *γ*_max_ = 10 agents show a reproductive disadvantage, intuitively because agents with larger *γ*_4_ tend to leave *S*_4_ resource levels (with a relatively high birth rate) more often and produce less offspring, while later in the evolutionary process this disadvantage is overcome by *γ*_4_ = *γ*_max_ agents in low densities as they eventually find enough available *S*_5_ sites to overcome their initial disadvantage. However, as density increases, other agents can quickly fill up *S*_5_ sites and leave little opportunities for *γ*_4_ = *γ*_*max*_ agents to overcome that initial disadvantage. Note that *b*_5_ = 5 is used in these numerical solutions so that 〈*γ*_4_(*t*)〉 converges to 10 at low densities.

Unlike the low-density model, we cannot tractably derive an effective objective function similar to Λ_*γ*_ in the high-density regime—in particular, it is not clear whether there exists a function *C*(*γ*) of a single phenotype vector *γ* whose optima predict the most competitive phenotypes. We discuss this issue further in Evolution as an optimization or learning process. Even if no such cost function for the phenotypes exists, one might wonder whether the dynamics for the (expected) population sizes can be written as a gradient descent of some cost function, 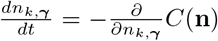. However, one can show that the vector field defined by the right hand side of Eq. (2) is not conservative, and therefore cannot be written as the gradient of an objective function *C*(**n**). The evolution of the population sizes therefore cannot be interpreted as optimization problem, at least using gradient descent dynamics.

## DISCUSSION

### Summary and interpretation of results

In this work we have used a population dynamics approach to explore the evolution of the innate foraging behaviors of a population of organisms exploring an environment with randomly distributed non-consumable resources. These organisms have limited sensory capabilities and no memory, and must take actions to either continue exploiting the resources at their current location or leave in search of locations with more resources (i.e., more favorable to reproduction). The phenotype of the organisms comprises the rates at which the agents attempt to relocate. While an “intuitive” prediction—the agents should leave low-resource sites as quickly as possible and not move from the highest resources sites—is indeed correct when the population density is low and high resource sites are readily available or have high enough birth rates to compensate their scarcity, this intuitive prediction becomes incorrect when the difficulty of finding high resource sites increases due to intrinsic scarcity or occupation by other organisms in high-density populations. In particular, as the initial population density varies, we not only observe a change in the dominant phenotype at long times, but we also observe long-lived periods of phenotype coexistence.

We have used a combination of stochastic agent-based simulations and a simplified mean-field model to gain insight into the basic phenomenology of the evolutionary dynamics. The resulting dynamics of the average population sizes are not typically interpretable as gradient descent-like learning dynamics, as might be obtained from a normative approach in which the organism phenotypes are predicted by optimizing a hypothetical objective function. Similarly, there is no guarantee that one can in general derive a function of the phenotype parameters that predicts the optimal phenotype—or distribution of phenotypes—without running the dynamics of the model.

We will discuss some of these points in detail in the following subsections, along with detailed remarks on Comparisons to related work and Limitations of the study, and we close with a brief discussion of Future directions.

#### Comparison of the agent-based and mean-field models

Overall, the outcomes of the full stochastic agent-based model and the minimal 5-state mean-field model we used to gain further insight were in qualitative agreement. This is often the case in statistical physics models; for example, mean-field approximations of Ising models of magnetism correctly predict the qualitative features of the phase diagram of the model, but overestimate the critical temperature at which a transition from zero to non-zero net magnetization of the material occurs [24], similar to the overestimate of the critical birth rate for the transition shown in Fig. 3. This discrepancy may also be due to the well-mixed assumption made during the reduction of the full 2*d* model to the 5-state model, discussed in the next paragraph. Despite some quantitative discrepancies in predicting the values of the critical birth rates *B*_5*c*_(*c*), the low-density mean-field model accurately predicts the transition boundary from 〈*γ*_4_〉 ≈ 10 to 〈*γ*_4_〉 ≈ 0 as a function of the resource scarcity *c*, as shown in Fig. 3.

The primary source of quantitative discrepancies between stochastic models and mean-field models is the neglect of fluctuations and the necessity of choosing a carrying capacity that takes continuous inputs (which gives rise to the factor 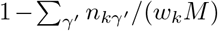 in Eq. (2). In many mean-field approximations of master equation models, upon which our 5-state model is based, the variance of these fluctuations eventually invalidate the mean field approximation at long times or when the mean population sizes are small. As discussed in Limitations of the study, we avoid these causes of mean-field breakdown by using a periodic catastrophe to reduce the population size to a fixed level, rather than stochastic death. Additional discrepancies are caused by the reduction from the mean-field approximation for the full 2*d* environment to the 5-state model (see Numerical solution and analysis of the 5-state mean-field model for details). Some of the discrepancy in Fig. 3 may also reflect the effects of population density, particularly in environments with scarce resources, which is outside of the regime of validity of the low-density model used to predict the transition boundary in Fig. 3B.

The largest qualitative discrepancy between the stochastic simulations and the mean-field model is the timescale of the evolutionary dynamics. Saturation of the hopping rates in the stochastic model appears to take ∼30 times longer than in the mean-field model, despite the birth rates, hopping rates, and catastrophe period parameters being nearly the same across the models. We believe this discrepancy is due to the well-mixing assumption of the mean-field model, which posits that the spatial variability of agents throughout the environment is negligible, allowing our reduction to the 5-state model. This conjecture is supported by direct simulations of a 5-state agent-based model, which shows a convergence time similar to the mean-field model predictions (see Supplementary Fig. 3). We therefore expect that in a mean-field model that takes into account spatial variation of agents and resources we would get convergence times similar to the full 2*d* agent-based simulations.

#### Metastability of phenotype coexistence

In both our stochastic simulations and mean-field model we observed long-lived coexistence of different phenotypes (Fig. 4 and 5). In the low-density limit of the mean-field model we could show analytically that this coexistence is almost-surely transient: only the phenotypes with the largest eigenvalues Λ_*γ*_ will survive in the limit of infinitely many catastrophes, *ℓT* → ∞. The exceptions in which coexistence is possible in this long time limit are fine-tuned conditions under which multiple phenotypes have the exact same values of Λ_*γ*_. Therefore, outside of this special fine-tuned condition, at low-density coexistence is *metastable* in the sense that phenotypes may coexist for long durations of time but the ultimate fixed point is survival of only a single phenotype (we remind the reader that extinction is not possible in our setup; in practice the fate of any model with a finite number of agents and stochastic death is extinction [25]). In the stochastic agent-based model the reproduction noise prevents any one phenotype from being the only survivor, but the distribution of phenotypes will be narrow, and long-lived coexistence is represented by a wide distribution of phenotype that persists across a large number of catastrophes. In both the mean-field model and the agent-based simulations the timescale of this phenotype coexistence depends strongly on the value of *b*_5_: when *b*_5_ ≈ *B*_5*c*_ the distribution of *γ*_4_ remains wide for a long time, while the distribution quickly converges when *b*_5_ ≪ *B*_5*c*_ or *b*_5_ ≫ *B*_5*c*_. This long-lived phenotype coexistence is again somewhat fine-tuned, requiring *b*_5_ to be close to *B*_5*c*_.

In the high density regime (Eq. (2)), on the other hand, we observed a range of densities for which multiple phenotypes coexist for long times and do not show clear signs of a trend towards a single dominant phenotype. This form of phenotype coexistence appears more robust—requiring less fine-tuning of the initial density—than the coexistence observed in the low-density regime when the birth rate *b*_5_ is close to the critical value *B*_5*c*_. However, it is difficult, if not analytically intractable, to determine whether the phenotype distribution would ultimately converge to a single dominant phenotype if run for a much longer time, or if there is a broader range of parameter space for which coexistence occurs in the infinite time limit. Inspection of the average hopping rate dynamics 〈*γ*_*k*_ (*t*)〉 in the mean-field model (Fig. 4F) seems to suggest there will be fine-tuned values of the initial density *N*_0_/*M* at which two or more phenotypes may coexist in the long-time limit, but deviations from these values result in a single surviving phenotype in the long time limit, rendering coexistence of phenotypes in the high-density regime similar to low-density coexistence near *b*_5_ ≈ *B*_5*c*_. However, because the mean-field model does not include reproduction noise, it is possible that the reproduction noise could stabilize coexistence states, preventing any one phenotype from dominating at long enough times in the full stochastic model—the trends of 〈*γ*_4_(*t*)〉 in Fig. 4A-B could be reflecting such a behavior.

#### Evolution as an optimization or learning process

In this work we investigated the evolution of innate foraging behaviors using evolutionary population dynamics [4], in contrast to the more common normative approaches in which evolution is assumed to act like an optimization process that extremizes some objective function subject to constraints [6, 26, 27]. One of our motivations for doing so was to better understand conditions under which we might expect such normative approaches to be mimicked naturally by the population dynamics, either through the population dynamics being interpretable as gradient-descent dynamics or by deriving an effective “fitness function” that predicts the dominant phenotype, or conditions under which typical normative approaches might need to consider “multi-agent” interactions to properly predict emergent phenotypes, a growing trend in deep reinforcement learning [28]. While normative methods have proven valuable for developing insight into biological structure in many areas of application [12, 17], conditions under which evolutionary dynamics can be expected to optimize abstract quantities such as “information” have not been extensively explored, and our goal was to gain insight into such limitations with a simple population dynamics model.

We found that at low initial population densities we could effectively capture the evolution of the agent phenotypes with a mean-field model that neglects interactions between different agents, and hence different phenotypes. Effectively, agents of different phenotypes only interact through the periodic catastrophe we use in our model to drive phenotype selection. In this regime it turns out that the overall growth rates of each phenotype are independent, and the catastrophes ultimately select for the phenotype with the largest growth rate. Thus, in this regime the maximum eigenvalue of the linear dynamics serves as an effective fitness function for the population that predicts the dominant phenotype at long times—this result is non-trivial, because *a priori* it was not obvious that the catastrophes, which nonlinearly couple the phenotypes, would not change the predictions of the linear eigenvalues. We might therefore hypothesize that organisms that evolved under low-density conditions in which individuals were not often competing for resources or space may be more amenable to normative approaches that aim to predict the optimal phenotype through the use of objective functions of only a single representative agent. This said, in a more realistic stochastic model with asynchronous death rather than a synchronous selecting catastrophe, there is no mechanism (other than extinction) for keeping the population size from eventually growing large enough to invalidate the low-density approximation. In an environment with a very large carrying capacity the low-density approximation may still be valid on the relevant evolutionary timescales, and the eigenvalues (growth rates) of such a mean-field model could still predict the phenotype that comprises the largest fraction of the population.

When the population density is sufficiently high that interactions between agents cannot be neglected (whether in our model with catastrophes or a model with stochastic death), then interactions between individuals can alter which phenotypes dominate the population at long times, or lead to long-lived coexistence, and there does not appear to be an obvious single-agent objective function that can be derived to predict the outcomes observed in our simulations or mean-field model. Moreover, the dynamics of the expected numbers of agents do not straightforwardly follow gradient-descent dynamics. This suggests it may not be possible to derive a function of a single representative phenotype parameter to exist in the interacting regime. However, this does not preclude the possibility of using machine-learning approaches to try and infer an effective objective function of just a single phenotype vector *γ*, though such a cost function may not be interpretable in terms of the actual evolutionary dynamics. A danger with this sort of approach is that one must be careful not to “overfit” proposed cost functions to observations. In principle, one can construct or infer a cost function to fit any observation, so the utility of a proposed cost function lies in how well it generalizes to other scenarios that it was not initially designed for [18]. Without imposing that an objective function and its associated constraints must generalize across novel conditions it was not constructed for, optimality of organisms in their environment becomes a tautology [4].

While the conditions under which evolution acts as an optimization process are not generally clear, and may not be valid in some circumstances [20, 29–31], evolution can perhaps still be interpreted as a kind of learning process. While the individual agents in our model do not possess any ability to learn, the population effectively “learns” the structure of the environment over the course of its evolution: the phenotype structure of the population after a long time depends on the structure of the environment and the dynamics of the agents. Even though there is not necessarily any well-defined fitness or objective function being optimized, the adaptation of the population to the environment displays some of the hallmarks of learning.

In this work we have only considered the action of evolutionary selection on certain aspects of an organisms’ innate behaviors, and have not allowed for the possibility of within-life adaptations that occur through development or decision-making behaviors. Evolution does not act directly on within-life adaptations—e.g., evolution could determine the overall structure of a species’ nervous system, but would not select for the synaptic weights between neurons. The synaptic weights reflect the experience of an organism and are ‘learned’ in the course of the individual’s early development. However, while evolution does not act directly on features like synaptic weights, it would act on the developmental and homeostatic rules that select those weights, and in this sense evolution could be interpreted as a kind of meta-learning that learns learning rules. Several lines of work have already begun investigating this kind of meta-learning using normative approaches, defining objective functions that select for learning rules rather than parameters of an agents’ structure or behavior [32–35].

### Comparisons to related work

Research studying the evolution of innate behaviors have focused on the interplay of the innate behavior evolution and learning within an individual’s lifetime. The two forms of behavior are found to be intertwined and share the same or similar neural circuits and evolutionary histories [36, 37]. A recent study hypothesised that innate behaviors and learned behaviors may have both evolved through epigenetic mechanisms, and thus studying the evolution of innate behaviors can enhance our understanding of learned behaviors and their physical underpinnings [38]. This said, the timescales separating the evolution of innate behaviors and within-life learning pose a challenge for exploring this co-evolution through the agent-based models and their mean-field approximations like we have considered here in this work.

Several lines of work have used population dynamics approaches based on master equation models to study the evolution of species under various ecological situations [25, 39], including competition for resources [40–43], prey and predation [44–47], evolutionary arms races [48, 49], and more. Our model follows this methodology, and may be viewed as a generalization of logistic population growth to a spatial environment with multiple possible phenotypes competing for the same limited resources.

The fact that our model considers only organisms of the same trophic level and does not include a hierarchy of organism types (i.e., similar organisms compete for a common pool of resources, as opposed to, e.g., predator and prey dynamics) is similar to the context of the neutral theory of biodiversity [42, 50–52]. The neutral theory of biodiversity assumes that the relative abundances of different species occupying the same environment is not necessarily due to relative evolutionary advantages or disadvantages, but potentially simply due to stochasticity of birth-death dynamics. That is, neutral theory considers it possible that differences between species may be neutral in terms of their relative competitiveness in an environment, and the species that dominates is due to “luck.” While the relative merits of the neutral theory of biodiversity versus niche-based theories have been a subject of debate in ecology [42], neutral theories have proved useful as minimal models in the vein we have presented our model here, and in general neutral models have been successful in explaining some observed trends in experimental data, although a more complete picture of ecological dynamics is thought to require non-neutral effects [40, 50]. In our model the different phenotypes have no intrinsic differences in terms of their raw birth rates—the birth rates are solely a function of the resource level at the agents’ location—so in this sense our population is neutral. However, phenotypes in our model do have emergent advantages over one another because some phenotypes are certainly more successful than others at finding high resource sites and producing more offspring, and thereby do possess selective advantages in our model. Our mean-field model in particular demonstrates that some phenotypes have clear advantages over others, and in the low-density limit we can derive the *fine-tuned* conditions under which different phenotypes can coexist in the long-time limit. As discussed in Metastability of phenotype coexistence, it appears that the stochastic agent-based model may have slightly more robust phenotype coexistence, possibly reflecting a mechanism of stochasticity-induced emergent neutrality between similar phenotypes for extended periods of time.

Beyond master-equation based models, there are many other studies that take normative approaches to understanding the behaviors and structures of organisms performing similar foraging tasks. Work in this vein may take several forms of abstraction, ranging from models in which the quantities to be optimized are directly relevant quantities—such as expected numbers of offspring [5] or the expected time it takes organisms to find a stockpile of food [6, 7]—to more abstract “rewards” such as the information an organism has extracted from its environment [11, 17, 27, 53, 54]. In some cases these normative models consider just a single representative agent and its associated parameters that determine its dynamics, and hence phenotype. More recent work has considered the dynamics of multiple agents through the lens of evolutionary game theory [17, 55–58], sometimes taking the limit of a continuous population density, effectively reducing the individual agent-based dynamics to a coarse-grained statistical description [27]. Many similar approaches are being developed in machine learning applications [28]. In some of these cases, such multi-agent cost functions are optimized using evolution-inspired “genetic algorithms” [59], in which the parameters of a population are remixed and inherited by the next generation after a certain amount of simulation time (i.e., there are typically no mechanistic birth-death events as in the model considered in this work).

Recently, multiple independent lines of work have begun formulating Bayesian approaches to bridge mechanistic and normative approaches to understanding biological structure [6, 9, 10]. Park and Pillow [9], M-lynarski *et al.* [10] are more suited to advancing our understanding of within-life adaptation and learning. Of particular relevance to our work presented here, Kilpatrick *et al.* [6, 7] extended classical optimal foraging theory with Bayesian updating to investigate optimal patch foraging strategies in both non-depleting and depleting patch environments. Their approach identifies different thresholding strategies for foragers in different environments and how the strategies are affected by forager’s uncertainty of the environment. Notably, the objective “minimize time to arrive and remain in highest yielding patch” is used as a premise when the forager is in the non-depleting patch environment, while in our work this “objective” emerges from the underlying birth-death dynamics.

As investigation of biological evolution from this multitude of approaches—mechanistic, normative or game theoretic, and Bayesian—continues to develop, it would be interesting for future studies to directly investigate the alignment of the outcomes of agent-based birth-death dynamics and normative approaches in such multi-agent settings, especially as increases in computational efficiency allow for simulation and investigation of increasingly complex agents.

### Limitations of the study

As with any model, there are many aspects of the phenomena we seek to study that cannot be fully captured by our approach or assumptions, and warrant discussion. We focus on discussing how some of the important model simplifications could potentially be relaxed, and consider possible implications for empirical data at the end of this section.

#### Disparity between evolutionary timescales and individual agent lifetimes

One of the most challenging aspects of modeling the evolutionary process is that the actions of individual organisms take place on time scales orders of magnitude shorter than the timescales over which significant phenotypic change occurs. This discrepancy is even more extreme for timescales over which accumulated mutations and reproduction result in radiations of new species. This separation of timescales presents an obstacle to modeling the evolution of complex behaviors and structures, motivating our choice of modeling agents with a very simple repertoire of available actions. Even relatively “simple” organisms like *C. Elegans* are capable of much more complicated behaviors than simply relocating based on the conditions at the organism’s current location [22]. While it is possible to consider models of more complex agents, for example with internal states that process environmental information, accounting for both the fast internal timescales and long evolutionary timescales in a single model is a significant challenge. Other modeling strategies for bridging these timescales will be necessary for the foreseeable future, such as the timescale separation approximation of Mann [27].

#### Selection by catastrophes versus stochastic death

We have implemented selection in our model by an unorthodox choice (in population dynamics models) of implementing a periodically occurring catastrophe that reduces the population down to a fixed size, rather than the more common and realistic scenario of having stochastic death in which agents randomly perish according to an asynchronous death rate (or set of death rates), rather than on a global clock. This said, this method of removing agents is similar to approaches used in evolutionary game theoretic models [17, 55–58]. The reasons for this modeling choice in our population dynamics approach are largely technical: in any stochastic model with a finite population size and stochastic death, the ultimate fate of the population is always extinction. While this may not be a problem in practice for simulations, as the extinction time can be quite large [25] and only a few simulations might terminate in extinction, it complicates analytic analyses. In particular, the mean-field approximation in a model with stochastic death will always predict that the extinction fixed point is unstable and the non-zero population size is stable, which is qualitatively invalidated at long times: the variance of the stochastic process grows with time, owing to the fact that eventually a large enough fluctuation will occur to drive the system to the extinction fixed point, from which it cannot escape. By only allowing agents to perish via a periodically occurring catastrophe that resets the population size to the initial number of agents we allow our model to remain close to the mean-field regime throughout the entirety of the simulation.

#### Sexual versus asexual reproduction

We considered only asexual reproduction in our model for simplicity, in which the phenotypes inherited by offspring are noisy copies of the parent’s phenotype (or direct copies in the mean-field model). One could introduce sexual reproduction into the model by allowing agents at adjacent sites to mate and produce an offspring onto one of the unoccupied sites adjacent to the parent that gives birth, with the child’s phenotype determined by some combination of the parents’ phenotypes.

The most significant impact that adding sexual reproduction would have on our results is the complication of the low-density approximation that we used to derive the effective cost function Λ_*γ*_. Because sexual reproduction requires interactions between individuals, a mean-field model that includes sexual reproduction is necessarily nonlinear. As a consequence, deriving an effective cost function like Λ_*γ*_ in this case is not straightforward, similar to the high density regime studied in this work. This could suggest that single-agent normative models may be most applicable not simply to organisms that evolved in low density conditions, but also organisms that reproduce primarily through asexual reproduction—or perhaps organisms that reproduction via external fertilization, such as many aquatic animals [60–63]. However, a more thorough analysis of phenotype dynamics in populations with sexual selection is required to test these possibilities.

#### Complexity of agents and environment in population dynamics models

The organisms we have considered in this work are rather simple in terms of their ability to process sensory information and take actions: an organism can only detect the resource level at its current location and move to a neighboring location at a fixed attempt rate. The agents have no memory, nor can they change their behavorial strategies, capabilities that even “simple” organisms like *C. Elegans* have been found to display after short periods of local exploration [22].

Even within the restricted repertoire of sensory processing and behavioral actions, we consider only relatively low-dimensional phenotype spaces, which in turn imposes a restrictions on the environmental structure: keeping the phenotype space constrained to 5 components requires we have only 5 resource levels. We could in principle have more resource levels while retaining a low-dimensional phenotype space by defining a function *γ*(*s*) parametrized by a small set of abstract parameters, but the choice of this functional form imposes additional structure, which could result in different competitive behavioral strategies for different choices. We therefore opted for the more parsimonious choice of a simpler environmental structure with a 1-to-1 correspondence with the (directly interpretable) hopping rate parameters.

Reasons for the simplicity of the agents include the issue of timescales discussed above and the combinatorial rate at which the phenotype space increases as more complex behaviors are permitted. For example, if agents’ hopping rates were functions of the agents’ current location and a potential target location, *γ*(*S*_*i*_, *S*_*j*_), then even for our simple 5-level environment there would be 5^2^ = 25 possible phenotype components corresponding to the 25 possible resource level changes *S*_*j*_ → *S*_*i*_, an exponential increase in the dimension of the phenotype space. Implementing even more complex agents will thus require even larger phenotype spaces. This is one of the largest challenges of modeling the evolution of biological organisms: while normative methods allow for complex, high-dimensional phenotype spaces (e.g., deep neural networks with millions of parameters), these are typically limited to a small number of agents. Investigating large populations of complex agents using the evolutionary birth-death dynamics of this work is computationally demanding, and hence we have focused on the more tractable extreme of relatively large populations of simple agents.

Finally, another limitation common to this evolutionary framework and normative frameworks is that they cannot generate phenotypes that were not pre-defined. For example, there is no ability for the agents in our model to evolve from only being able to detect the resource level at their current location to having the ability to detect the resource level at nearby sites as well. Phenomenologically, this can be interpreted as being due to very long timescales for beneficial mutations that lead to the radiation of new genotypes resulting in a larger phenotype space, compared to the timescales of the simulations. However, this interpretation can be relaxed and this problem can be partially overcome by allowing for a larger possible phenotype space than is initially present in the initial population.

#### Depleting versus non-depleting resources

In this study we have considered only non-depleting resources (also called “non-destructive resources”), such that the resource levels *S*_*k*_ are not modified by the dynamics of the organisms in the environment. This restricts the interpretation of these resource levels to “conditions favorable for reproduction,” such as temperature or shelter, or consumables that are so plentiful that consumption by agents does not appreciably change the resource level of a site. Consumption of resources with replacement (e.g., continued growth of food sources or rainfall) would allow for a feedback loop between organisms and the environment, as the actions of the organisms change the environment, which in turn can alter which behavioral phenotypes are most successful at finding sites with plentiful resources. In such a scenario the dominant phenotype could be constantly changing, unless the statistics of resource consumption of and agent dynamics achieve an equilibrium. Such a scenario could give rise to the evolutionary dynamics similar to those observed in populations of turtle ants,in which the head shapes of soldier ants have repeatedly reappeared through evolution [20], though the specific case of soldier ants likely involves additional environmental selective pressures that change with time, not just changing resource distributions. Modeling such systems from a normative perspective would require defining a fitness landscape that changes with time. Investigating these possibilities within frameworks like ours will be an interesting avenue for future work studying the evolution of organism behaviors and sensory capabilities.

#### Metabolic costs of reproduction and relocation

In our model we have not explicitly implemented any metabolic penalty for giving birth or hopping to a new location. The phenomenological effects of such metabolic costs are implicit in the resource level dependence of the birth rates and the imposition of maximum hopping rate *γ*_max_. A more complex agent-based model could give each agent its own metabolic budget. Agents would acquire energy from the environment—e.g., high resource sites would provide more energy per unit time than low resource sites, possibly through consumption of more plentiful resources—and then agents would spend that energy to give birth (incurring a large metabolic cost) or relocate, with the metabolic cost of moving depending on the speed (hopping rate) at which the agents attempts to move. In such a formulation the hopping rate would not need to have an intrinsic maximum per se, though there would be an effective maximum determined by the ability of the agents to acquire enough food to support the high metabolic costs of moving quickly.

A metabolic budget could be incorporated into a master equation model or mean-field model by treating the energy as an additional variable on par with the agent position and phenotype, i.e., by counting the number of agents at a particular location with a particular phenotype configuration and specified amount of energy.

We expect that in an environment with non-consumable resources that provide agents with a fixed amount of energy per unit time, with higher rates conferred by higher resource levels, one could choose the energy rates and metabolic costs of reproduction and relocation in such a way that the results of our current study are qualitatively reproduced.

#### Comparison to empirical data

Owing to many of the simplifications of our model discussed above, it is difficult to directly compare our results to empirical data. The most promising means of comparing our model to empirical data would be to design and perform an experiment that closely mimics the set-up of the model. For instance, one could imagine an experiment with a population of organisms that a) are capable of self-propelled motion, b) have some means of detecting local nutrient level/temperature/etc., c) have tools for genetic modification to create mutants representing an initial distribution the phenotype behaviors, and d) have relatively short reproduction times. These organisms would be placed into a controlled environment in which nutrients or reproductively favorable conditions are randomly distributed; consumable resources would need to be replenished at rates comparable to the rate of uptake by the organisms. The organisms would be given time to explore their environment and discover locations of high nutrient content, after which they are culled back to the initial population size, mimicking the catastrophe in our model. By using tracking software to monitor the organisms’ relocation rates out of each nutrient level, one could observe over a large number of cullings whether there is a change in the distribution of phenotyped behaviors over the course of the experiment.

This said, the implementation of such an experiment may not be pragmatic for the foreseeable future, and would likely be limited to single-celled organisms. A more promising avenue might be to compare the qualitative predictions of future extensions of this work modeling more complex organisms. For example, experimental studies have observed selection-induced genetic drift in a mixed population of wild-type mice and mice with a mutation that disrupts their circadian rhythm [64]. One could model such an experiment by incorporating some of the more complex features discussed above into a formalism like the one presented in this work, with additional modifications to introduce a a day/night cycle that interacts with agents’ internal circadian rhythms. It would be of great interest to model similar scenarios that investigate the relative reproductive success of organisms with different perturbations to their sensory modalities, to see if the outcomes can be assessed using various normative predictions commonly used in neuroscience.

### Future directions

In this first work we have focused on agents with relatively simple sensory capabilities and behaviors in order to tractably compare agent-based simulations with mean-field approximations and calculate an effective fitness function in a tractable limit of the mean-field model. Two main avenues for future investigations of the models that we have not attempted to elucidate here include i) performing more advanced mathematical analyses of master equation models, for example by investigating the effect of stochastic fluctuations on the stability of phenotype coexistence in more detail, and ii) pushing the boundaries of agent and environment complexity, along with implementing other more realistic features of evolutionary dynamics.

Investigating the effects of stochasticity requires tools to go beyond the mean-field models introduced here. Several tools exist in the mathematical and statistical physics literature for this purpose, such as the Wentzel–Kramers–Brillouin (WKB) approximation [25, 39] or field-theoretic formulations of the master equation model [44–46, 65]. Because these cases explicitly deal with beyond-mean-field effects it would be more natural to use stochastic death in this setting, rather than the periodic catastrophe introduced in this work to control the efficacy of the mean-field approximation. For example, the WKB approximation could allow for a direct calculate of the large deviation functions determining the extinction transition.

In terms of increasing organism complexity, a long-term goal of this line of work would be to endow agents with “brains”—small neural networks that take in some sensory information from the environment and then determine how the agent will attempt to move. In such a context evolution would act on the structure of the brain, similar to the case of Braitenberg vehicles [66, 67] (as opposed to determining the synaptic weights in neural network models such as in [68]). In practice, this will be a challenging direction due to the large phenotype space and the separation of timescales involved in sensory processing and evolution. A more tractable approach would be to endow the organisms with a simple “receptive field” that detects sensory information in its immediate environment, down to a modest modification of the current model that allows for agents to detect resource levels at neighboring sites in addition to its current location. Straightforward increases in environmental complexity include adding spatial correlations between resources levels at different locations, as well as introducing sensory signals distributed through the environment that are correlated with resource levels that agents cannot detect directly. Modeling the impact of consumable resources is also of ecological importance, and could be implemented in our models, e.g., by introducing a non-motile “species” at each site that is consumed by agents and replenished at a fixed rate. In this way one could also introduce more realistic metabolic limitations: actions such as giving birth or relocating could incur energy costs that must be replenished by consuming food in the environment. This is one way we could formally remove the artificial restriction of a maximum hopping rate *γ*_max_, allowing the organisms to move as fast as possible between sites if they can acquire the necessary energy to maintain such a strategy.

Finally, one of the goals of this work was to attempt to derive functions that could serve as effective cost functions in normative approaches to modeling the biological structures and behavioral strategies that emerge from the more fundamental evolutionary dynamics. It would be interesting to perform a more thorough comparison of the predictions of birth-death dynamics and normative approaches to identify regimes where the predictions diverge, especially in approaches in which the objective functions are abstract quantities such as information-theoretic quantities related to random variables in an organism’s environment [9–11]. Doing so will provide valuable insight into the relationships between fundamental evolutionary processes and the more parsimonious normative approaches useful for modeling complex organisms.

## METHODS

### Details of the agent-based stochastic simulations

In the agent-based stochastic simulation, we use an *L* × *L* lattice with periodic boundary conditions, with a standard value of *L* = 128. The resource level at each location is drawn from a discrete uniform distribution with 5 possible levels, labeled *S*_1_, *S*_2_, *S*_3_, *S*_4_, and *S*_5_. Each agent has a phenotype vector *γ* = (*γ*_1_, *γ*_2_, *γ*_3_, *γ*_4_, *γ*_5_) where the *i*^*th*^ component corresponds to the hopping or relocation rate when the agent is at a site with resource level *S*_*i*_. The birth rate of agents at spatial coordinate x within the environment is a function of the resource level at that location, *b*(*s*_x_) = *b*(*S*_*i*_) ≡ *b*_*i*_ if *s*_x_= *S*_*i*_. Initially, 500 agents are introduced to the system, each of which has a random initial location and random initial phenotype vector with each component ranging from *γ*_min_ = 0 to *γ*_max_ = 10. This bound is set considering that the relocation rate cannot go negative and there are metabolic costs that constrain agents from relocating too frequently. An offspring inherits its parent’s phenotype vector with Gaussian noise added to each component. The noise term follows 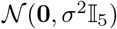, with *σ* = 0.05 (arbitrary units) in all the stochastic simulations throughout this study. If the noise would push a relocation rate component outside of the range [*γ*_min_, *γ*_max_] then the component is set to the closest boundary value. Each location can hold at most 1 agent at any given time and thus the reproduction process has to leave the offspring at one of the vertical or horizontal neighboring locations in practice—if the agent attempts to give birth into an occupied site then this reproduction attempt is considered as unsuccessful. Similarly, hopping may only occur from an agent’s position to one of the four adjacent sites, with the hop being unsuccessful if the target site is occupied.

Importantly, a system-wide periodic random culling is applied to the system after the stochastic simulation after a duration of *T* = 0.3 (arbitrary time units), which reduces the number of agents back to its initial state. This periodic catastrophe removes agents in a random manner without distinguishing its phenotype or location, functioning as the selection in the evolutionary dynamics. We expect that the most competitive phenotypes will be selected out after a series of growth and catastrophe events. A full iteration of the simulation consists of stochastic growth and relocation, as well as a random culling at the end. The next iteration starts off with the survived agents from the last iteration as the new initial condition.

As the carrying capacity of each location is set to 1, the 128 × 128 2*d* lattice can hold at most 16384 agents. For a typical run, in 0.3 units of simulation time the number of agents grows from 500 to around 850 before the random culling brings the number of agents back to 500. The initial density of agents is defined as initial number of agents divided by total possible number of agents in the system, which is 500/16384 ≈ 3.1% for this standard parameter set. When investigating the high density regime, we kept the number of agents to be 500 in all the initial conditions and reduced the lattice size from 128 × 128 to smaller size to achieve higher initial densities.

To reduce the potential variability from stochastic simulations, we run 20 independent simulations with the same set of model parameters and resource background for all the agent-based stochastic simulations in this study. Then the averaged phenotype components cross all trials and their standard deviation were calculated as the solid lines and shaded areas in Fig. 2E and Fig. 4A-D.

The stochastic agent-based model was simulated in the Python programming language using a direct method Gillespie algorithm [23]. The Gillespie algorithm computes the total rates of the possible events that may take place in the system to determine when and what the next “action” is. Here “action” refers to either birth or relocating of an agent in the system. The total rate is the sum of the birth and relocation rates of all the agents in the system; the algorithm first determines the time of the next action and then probabilistically chooses one of the available actions based on the relative rate of each action compared to the total rate. That is, at each step, the time of the next birth or relocation event is drawn from an exponential distribution with mean 1/total rate. The probability that the next event is agent *A* giving birth or relocating is *A*^′^s birth rate/total rate or *A*^′^s relocation rate/total rate. The target location of the birth or relocation event is then randomly chosen from one of the four possible neighboring locations of the agent’s current position. If the chosen target location is already occupied, the birth or relocation attempt is considered unsuccessful. Note that in this implementation of the algorithm, hopping or birth from site x to site **y** is a two-step process: the rates going into the total rate are just *b*(*s*_x_) and *γ*_x_, and *then* if one of these events is chosen to occur the target site is drawn with probability 1/2*d* = 1/4 in our two-dimensional environment. However, an alternative way we could have implemented these dynamics is via a one-step process in which we include the birth and hopping rates from site x to site **y** for all possible target sites as separate rates into the total rate (all of which happen to have the same numerical values). In this second possible implementation the birth and hopping rates must be scaled down by the number of target sites (2*d* = 4) in order to match the results of the two-step implementation. This technical detail is important when developing the mean-field model, as the one-step implementation corresponds more directly to the master equation formalism we introduce in Master equation model of the stochastic dynamics and derivation of the mean-field approximation, and scaling the birth and hopping rates is required to make the derived mean-field model match the agent-based simulations.

Multiple Python packages were used in the stochastic simulations, mean-field model analyses as well as data visualizations in this study, most notably, Numpy [69], Scipy [70], and Matplotlib [71]. In the stochastic simulations, each stochastic trajectory was run in series on a single CPU core. A computing server with two Intel^®^ Xeon^®^ Processor E5-2690 v4 CPU (14 cores for each CPU unit) was used to generate 20 stochastic trajectories for the same set of parameters in parallel with the aid of the Python Multiprocessing module. Typically, each run takes approximately 35 minutes for a system starting with 500 agents and *b*_5_ = 4 in Fig. 2. Higher *b*_5_ led to more agents being produced, resulting in a higher simulation load, and thus simulations with higher *b*_5_ value take longer to run. For example, the simulations with *b*_5_ = 16 depicted in Fig. 3 took over an hour to run.

### Master equation model of the stochastic dynamics and derivation of the mean-field approximation

#### Full model in a d-dimensional environment with random resources

Formalizing agent-based stochastic simulations as mathematical models that allow for tractable analytic analyses is often difficult [72]. Instead of keeping track of every agent individually, an equivalent description that lends itself to a standard class of mathematical models is one in which we only keep track of the number of agents with each phenotype at each spatial location in the environment. In order to write down such a model we need to make a few simplifications:

1. We discretize the phenotype space *γ* so that there are a finite number of discrete phenotypes and the number of agents with each phenotype can be enumerated. In principle, we could work with a continuous phenotype space and define a density of agents with each phenotype, but for practical numerical calculations using the mean-field model it is necessary to discretize the phenotypes.
2. We omit reproduction noise. In the mean-field model we will formally include all possible phenotypes in the initial conditions, so we do not need reproduction noise to introduce any phenotypes that were not initially present, as is the case in the agent-based simulations.

Now, let *n*_x,*γ*_ be the number of agents at location x with a particular phenotype *γ* (i.e., a particular choice of values for each of the relocation rate components) and denote by **n** the collection of all of these variables into a single vector. The probability that the state of the system has a particular configuration **n** at time *t*, *p*(*t*, **n**), is the solution to the system of differential equations

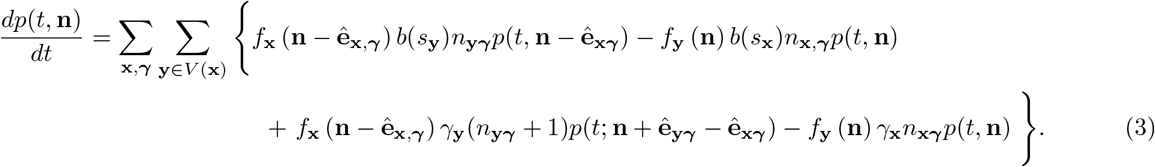

We have suppressed the conditional dependence of *p*(*t*, **n**) on the configuration of resource levels *s*_x_. The notation 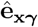 represents a vector with a 1 at the element labeled by (x, *γ*) and 0 elsewhere. The bare birth rate per capita at a site x is *b*(*s*_x_), explicitly denoting that the birth rate depends only on the resource level of the current location. We assume that each site has a maximum occupancy of a single agent, so all births occur via an agent at some site **y** birthing a new agent into an adjacent site x. The bare hopping rate out of a site **y** is denoted *γ*_**y**_; the value of the hopping rate is equal to the hopping agent’s value of the hopping rate at resource level *s*_**y**_. Both birth rates and hopping rates into a site x are modulated by a factor *f*_x_(**n**), which is 0 if site x is occupied and 1 if unoccupied. The possible target sites for hopping or birth by an agent at site x are identified by the notation **y** ∈ *V* (x), which means that site **y** is in the “vicinity” of x. By assumption, x ∉ *V* (x). For our 2*d* simulations, *V* (x) consists of the nearest neighbor sites **y** = x ± (1, 0)^*T*^ and **y** = x ± (0, 1)^*T*^.

The initial condition *p*(0, **n**) represents the initial configuration of the system. In many applications the initial condition is deterministic, but one can choose an initial distribution as well. In particular, we can ensure every phenotype is represented by choosing an initial distribution of *N*_0_ agents and assigning each agent’s phenotype and location by drawing from the possible phenotypes and sites with a uniform probability. This eliminates the need for reproduction noise in this model.

The dynamics of Eq. (3) run from *t* = 0 up to *t* = *T*, at which point the catastrophe occurs. *N*_0_ agents are randomly selected out of the agents alive at time *t* = *T* and retain their spatial position and phenotype as we eliminate the remaining agents from the environment. The distribution of remaining agents forms the initial distribution for the next iteration. This procedure repeats indefinitely, but we expect the distributions of the total number of agents of each phenotype to converge after a large number of catastrophes, though it may not be the case that the full spatial distributions *p*(0, **n**) and *p*(*T*, **n**) converge.

By summing Eq. (3) over all possible configurations **n**, changing dummy indices, and using the fact that *p*(*t*, **n**) = 0 if any *n*_x,*γ*_ < 0, one can show that the right hand side vanishes, as it must in order for probability to be conserved.

In practice, solving probability flow equations such as Eq. (3) is intractable except for very simple cases. Moreover, implementing the catastrophe at this level of description is challenging. A more common approach that lends itself to analytical and numerical analyses is to instead derive a system of equations for the moments of the distribution and make a mean-field approximation to close the system of equations (as moments are normally coupled to higher order moments). For example, the “mean-field approximation” corresponds to deriving a system of equations for the first moments of the distribution and assuming all higher order cumulants vanish, such that averages over functions of **n** can be replaced by those functions evaluated at the average 〈**n**〉. For Eq. (3) the mean-field system of equations is

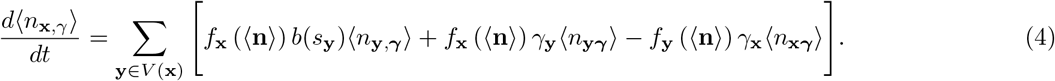

where we employed the mean-field approximation to replace products such as 〈*f*_x_(**n**)*n*_x,*γ*_) with the factored moments *f*_x_(〈**n**〉)〈*n*_x,*γ*_〉. Note that in order for this approximation to work with our modulation function *f*_x_(**n**) we must specify the values that this function *f*_x_ takes for continuous-valued inputs, as 〈**n**〉 will not be an integer. In this work we make the choice

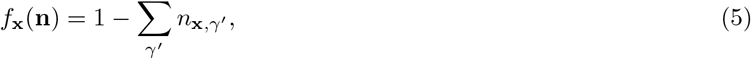

which satisfies the desired boundary conditions of *f*_x_ = 1 when the site is unoccupied and 0 when there is an agent at the site. This choice amounts to the assumption that 〈*f*_x_(**n**)*n*_x,*γ*_〉/〈*n*_x,*γ*_〉 varies approximately linearly with 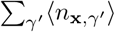. Although it appears that we should also impose a threshold to prevent this choice of *f*_x_(**n**) from becoming negative, this is unnecessary in practice as long as the initial conditions respect the occupancy of the system, as once *f*_x_ reaches 0 it blocks any further increases in occupancy that would drive it to be negative. This is true in both the full master equation model and the mean-field model (4).

These mean-field equations are still challenging to analyze due to the spatial variability of the system. We simplify our analysis further by deriving an approximate system for the total number of agents at sites of each resource level, detailed in the next section.

#### Reduction to a 5-state mean-field model

Rather than keeping track of the expected number of agents of each phenotype at every location in the environment we will instead focus only on the number of agents of each phenotype at sites of each resource level *S*_*k*_. The total expected number of agents at resource level *k* is

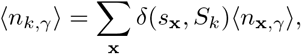

where *δ*(*s*_x_, *S*_*k*_) is an indicator function that is 1 if *s*_x_ = *S*_*k*_ and 0 otherwise. We will denote spatial sites by bold x and **y** indices while states will be denoted by italic *k* and *i* indices.

We can derive the system of equations for ⟨*n*_*k,γ*_⟩ by multiplying Eq. (4) by *δ*(*s*_x_, *S*_*k*_) and summing over x. The restricted sum over **y** ∈ *V* (x) poses a complication for this approach, and will require an approximation to be discussed below. We detail the calculation for each term of Eq. (4) in turn.

The left-hand side of Eq. (4) is trivial, and simply gives 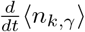, so we consider the birth rate term 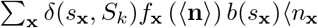 first. Note that we may write 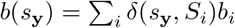, which essentially just says that *b*(*s*_**y**_) = *b*_*i*_ if *s*_**y**_ = *S*_*i*_. Proceeding,

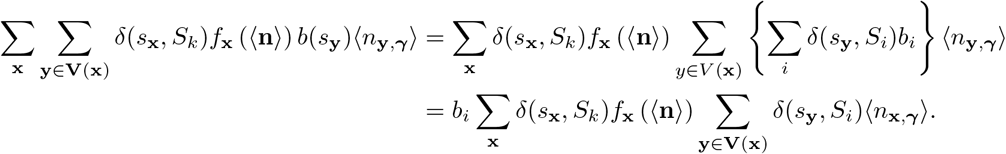

The sum over **y** depends on the current value of x, and so cannot be easily evaluated. One way to circumvent this problem would be to make the extreme approximation of allowing the target sites to include not just nearest neighbors but *all* sites in the environment (a usage sometimes used synonymously with “mean-field approximation,” but this is due to the fact that the mean-field approximation becomes exact on all-to-all graphs in many models). A slightly less drastic approximation is to assume that the environment is “well-mixed,” such that the sum over a local set of sites should just be a scaled down version of the sum over all sites. Specifically, we will assume

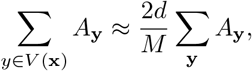

where the sum on the right-hand side is over all sites **y** and *A*_**y**_ is a general observable. The ratio 2*d/M* is the number of target sites (nearest-neighbors) in *d* dimensions to the total number of sites in the environment. For *A*_**y**_ = *δ*(*s*_**y**_, *S*_*i*_)〈*n*_**y**,*γ*_〉 we therefore obtain

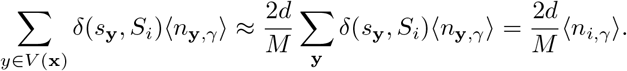

This approximation of the sum no longer depends on x, so we may now evaluate the sum over the modulation function *f*(〈**n**〉). For our choice of the modulation function we have

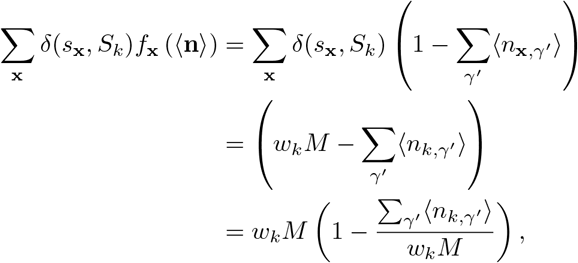

where 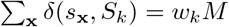 is the expected number of sites of resource level *k*, with *M* the total number of sites and *w*_*k*_ the frequency of resource level *k*. We assume the number of sites is large enough that the actual number of sites of each level is equal to its expected value. Thus, the birth rate term evaluates to

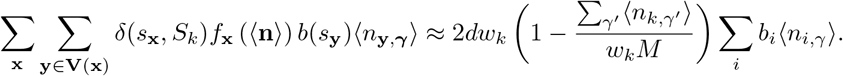

Next we consider the first hopping rate term. For this term it will be useful to use the identity 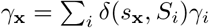, which like the similar identify for the birth rates just says that *γ*_x_ = *γ*_*i*_ if *s*_x_ = *S*_*i*_. Then,

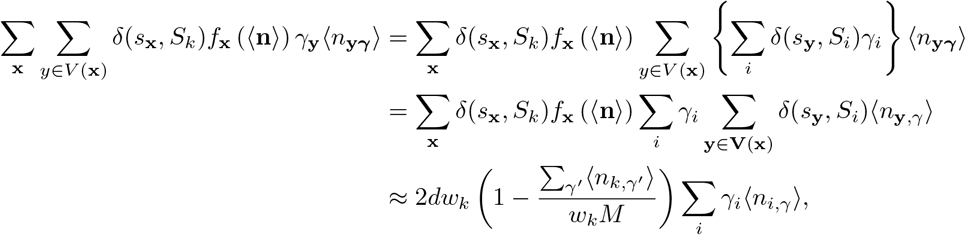

where we used the same approximations for the sums over **y** and x as in the approximation of the birth rate term. For the final term we have

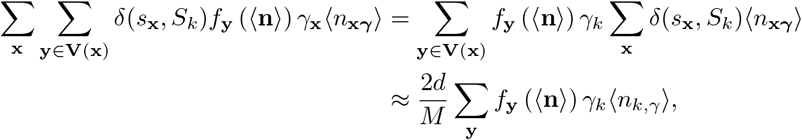

where the equality in the first line follows from using the fact that *δ*(*s*_x_, *S*_*k*_)*γ*_x_ = *δ*(*s*_x_, *S*_*k*_)*γ*_*k*_, and we used our approximation for the sum over nearest-neighbor sites as a fraction of the sum over all sites in going to the second line. For our choice of *f* the sum over **y** is

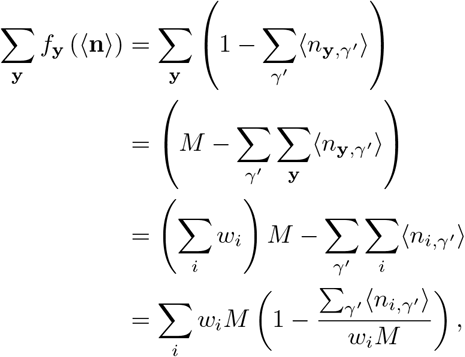

where we used the fact that 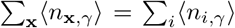 (the total number of agents obtained by summing over all sites must be equal to the total number of agents obtained by summing over all states) and the total frequency of each state must be 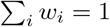. The final term of the mean-field equation is thus

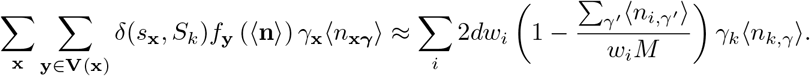

Putting everything together, the system of differential equations for our 5-state model is

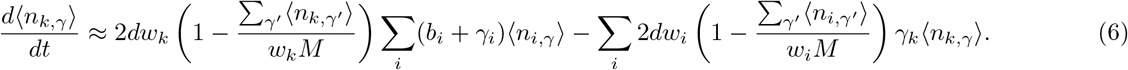

This is nearly our final result for the mean-field model. One additional correction is necessary to align it with our agent-based simulations.

#### Conditional-event correction and the low-density approximation

The 5-state model Eq. (6) describes the mean-field approximation to Eq. (3), which would correspond to a direct Gillespie simulation of the master equation (3) that treats all of the possible events at any moment on equal footing when calculating the total rate to determine the next event. For example, birth from site x to x ± (0, 1)^*T*^ and birth from x ± (1, 0)^*T*^ are all treated as separate events, even though they have the same rate (when the target sites are all unoccupied). The same is true for the different hopping directions.

However, as detailed in Details of the agent-based stochastic simulations, our implementation of the agent dynamics uses the fact that the birth and hopping rates only depend on an agent’s current location, not its target location, to calculate the total Gillespie rate using only a single instance of the birth or hopping rates. Then, *conditional on a birth or hopping event being determined to occur*, the target site is chosen randomly from one of the possible targets with equal probability.

These two implementations of the Gillespie dynamics are *not* equivalent. However, they are simply related. The conditional birth/hopping implementation essentially reduces the birth and hopping rates by a factor of the number of target sites. In a *d*-dimensional lattice this means we should rescale the rates *b*_*i*_ → *b*_*i*_/2*d* and *γ*_*i*_ → *γ*_*i*_/2*d* in Eq. (6), yielding the mean-field dynamics that correspond to our implementation of the stochastic simulations,

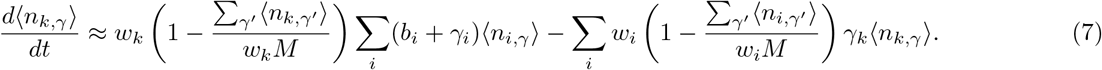

The mean-field equations given in the main text follow from this equation. In particular, the low-density approximation follows from this system of equations by assuming that 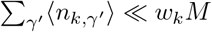, giving

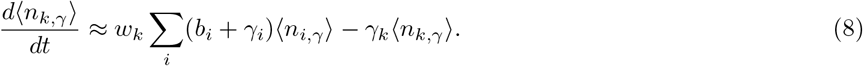

for each phenotype *γ*. As this is a linear equation, we may also write it in matrix form,

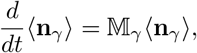

where 〈**n** _*γ*_〉 is a 5 × 1 vector of the expected number of agents of phenotype *γ* at each of the 5 resource levels and 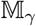 is the transition matrix involving the birth and hopping rates. This equation can be formally solved in terms of the matrix exponential: 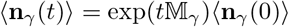, for each *γ*.

Note that although the phenotypes have become dynamically independent in this low-density approximation, the catastrophe still couples them together, as the fraction by which the population size is reduced depends on all of the phenotypes. We give our mathematical analysis of the effect of catastrophes in the next section.

#### Catastrophes in the mean-field model

We introduce catastrophes into the dynamics of Eq. (7) by running the mean field dynamics up to time *t* = *T*, at which point the catastrophe occurs and we reset the population to its initial size *N*_0_ by uniformly reducing the size of each phenotype:

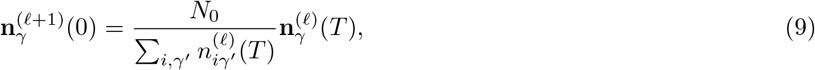

where *ℓ* indexes the number of catastrophes that have occurred and we have dropped the angled brackets for convenience. Though this equation holds for any density, in the low-density limit we can formally solve Eq. (8) and use this iterative map to derive the effective objective function Λ_*γ*_, the growth rate of the phenotypes, as we show below.

#### At low-density the most competitive phenotype has the largest maximum eigenvalue

We now prove that the most competitive phenotype in the reduced mean field model is that whose hopping rates maximize the largest eigenvalue of 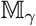, denoted Λ_*γ*_.

Our proof begins by using the explicit solution of Eq. (8) in terms of the matrix exponential, together with the reset condition Eq. (9), to relate the final population sizes just before the next catastrophe by a system of iterative maps:

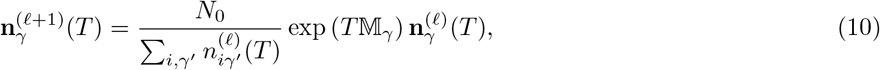

where 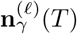 is the vector of population sizes at the end of the *ℓ*_th_ iteration (i.e., just prior to the *ℓ*_th_ catastrophe). As *ℓ* → ∞ this iterative map may converge to a fixed point, 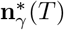, which is the stable distribution of the population in this dynamical system. The fixed point equation may be rewritten

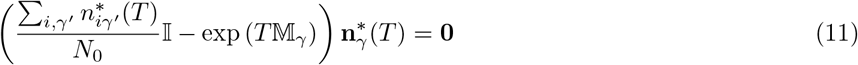

for each phenotype *γ*, and where 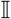 is the identity matrix. We expect the trivial solution 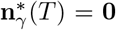 to be obtained for phenotypes that are outcompeted by other phenotypes, such that these phenotypes comprise smaller and smaller fractions of the total population, eventually vanishing as *ℓ* → ∞.

Because extinction is not possible in this model, at least one phenotype must achieve a non-zero population size at the fixed point. For a non-trivial fixed point to exist, the determinant of 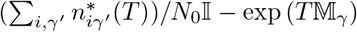 must vanish for some phenotypes *γ*. This implies that 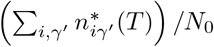 is an eigenvalue of the matrices 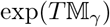 and 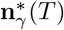 is the corresponding eigenvector. The eigenvalues and eigenvectors of 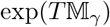 are related to the eigenvalues *λ*_*γ*_ and eigenvectors of the much simpler 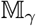, which can be efficiently computed ([73]). As empirically observed in Fig. 6, only one of the eigenvalues of 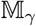 is positive; the rest are negative. As the self-consistency condition requires that 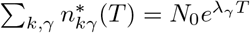, the only possible choice of eigenvector is that which corresponds to the largest, positive eigenvalue, which we denote Λ_*γ*_, as otherwise 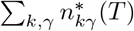 would be less than *N*_0_, which is not possible in this model. (Moreover, the eigenvectors with negative eigenvalues often have negative components, which does not make sense as a population count).

**FIG. 6.**
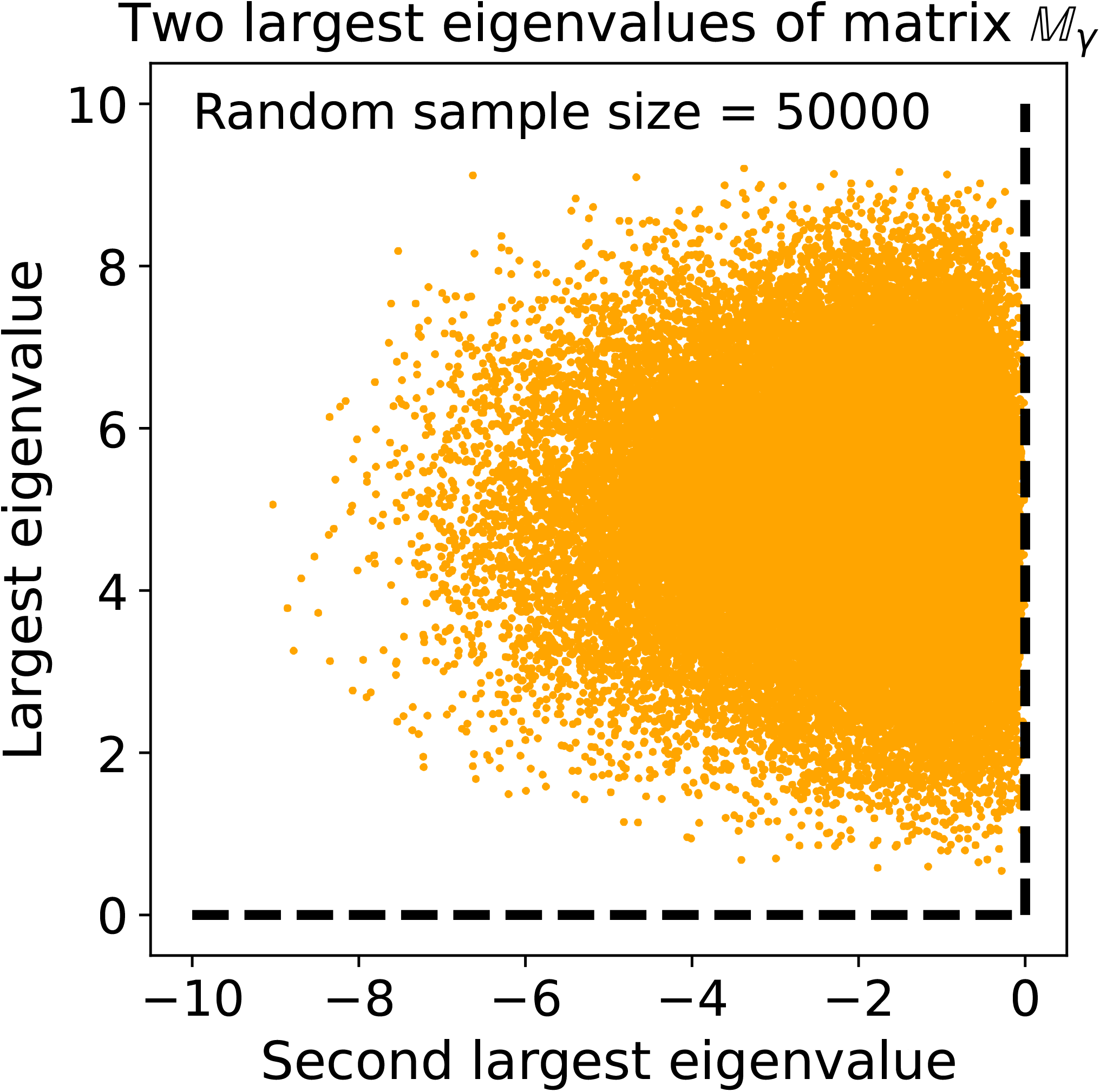
Empirical demonstration that the low-density coefficient matrix 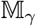 has only one positive eigenvalue. We numerically calculate the eigenvalues of the matrix 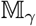 by randomly drawing all birth and relocation rates from a uniform distribution in [0, 10). Across 50000 samples shown, we find that only the largest eigenvalue of this matrix is positive and all the other eigenvalues are always negative.

Because we assume that all phenotypes are present in equal proportions in the mean field model, it follows that the phenotype with the largest overall maximum eigenvalue Λ_*γ*_ will be the sole survivor in the *ℓ* → ∞ limit. The fact that 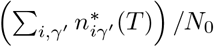, a single scalar number, must be an eigenvalue of 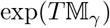 for *every γ* implies that the only way multiple phenotypes may coexist in the low-density limit is if their largest eigenvalues Λ_*γ*_ have the same value. In such a case the relative proportions of those phenotypes will depend on the overlap of **n** (0) with the eigenvector corresponding to Λ_*γ*_, which could be different for different phenotypes even if their eigenvalues are the same, leading to different relative proportions in the long time limit.

### Numerical solution and analysis of the 5-state mean-field model

With the simplified 5-state mean-field model we can evaluate the evolution of agents by numerically solving the system of differential equations in Eq. (7). To do so, we discretize the continuous phenotype space *γ* = (*γ*_1_, *γ*_2_, *γ*_3_, *γ*_4_, *γ*_5_) with *γ*_*i*_ ∈ [0, 10] into a space where each component can take discrete values from the set {0, 2, 4, 6, 8, 10}. This discretized phenotype space contains 6^5^ = 7776 possible phenotypes.

We numerically solve Eq. (8) with an ODE solver from Scipy package [70], using explicit Runge-Kutta method of order 5(4). In practice, we solve the full mean-field model Eq. (2) for the fully coupled phenotype system, even in the low density limit. The number of agents is reset following Eq. 9 after a duration of simulation time *T* = 0.3, repeated for 2000 iterations. A typical run takes around 2 hours to finish. Note that in all the numerical evaluations of the mean-field model, the birth rates are taken to be [0, 1, 2, 3, 5] except for Fig. 3 where the change of *b*_5_ is explicitly investigated. This slightly different choice of *b*_5_ = 5 in the mean-field model compared to the *b*_5_ = 4 in the stochastic model is to compensate the minor quantitative difference of the critical value of *b*_5_ at which the transition from 〈*γ*_4_〉 = *γ*_*max*_ to *γ*_*min*_ occurs, as shown in Fig. 3A, upper panel.

The mean phenotype and its standard deviation can be calculated from the fraction of agents with different phenotypes

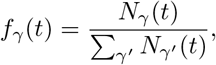

where 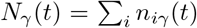 is the mean total number of agents with phenotype *γ*. Because there are no deaths in our model the numerator will never go to zero, and *f*_*γ*_ is therefore bounded between 0 and 1, so we can treat it as a probability and calculate the mean phenotype by

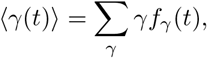

for each component of the vector *γ*. We similarly estimate the standard deviation by

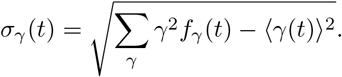

## ACKNOWLEDGMENTS

We thank Ken Dill, Piotr Sokół, Jiahao Hu, and Wenbin Lu for helpful discussions, and the Python community for building the tools used in this work.

## SUPPLEMENTARY INFORMATION

### Maximum eigenvalue as the effective fitness function

As discussed in the main text, the maximum eigenvalue behaves as an effective objective function and predicts the transition of *γ*_4_ when *b*_5_ is varied. It also explains the large variability of the *γ*_4_ component, as the maximum eigenvalue is insensitive to *γ*_4_ when *b*_5_ ≈ *B*_5*c*_, shown in Supplementary Fig. 1 below.

**FIG. 1.**
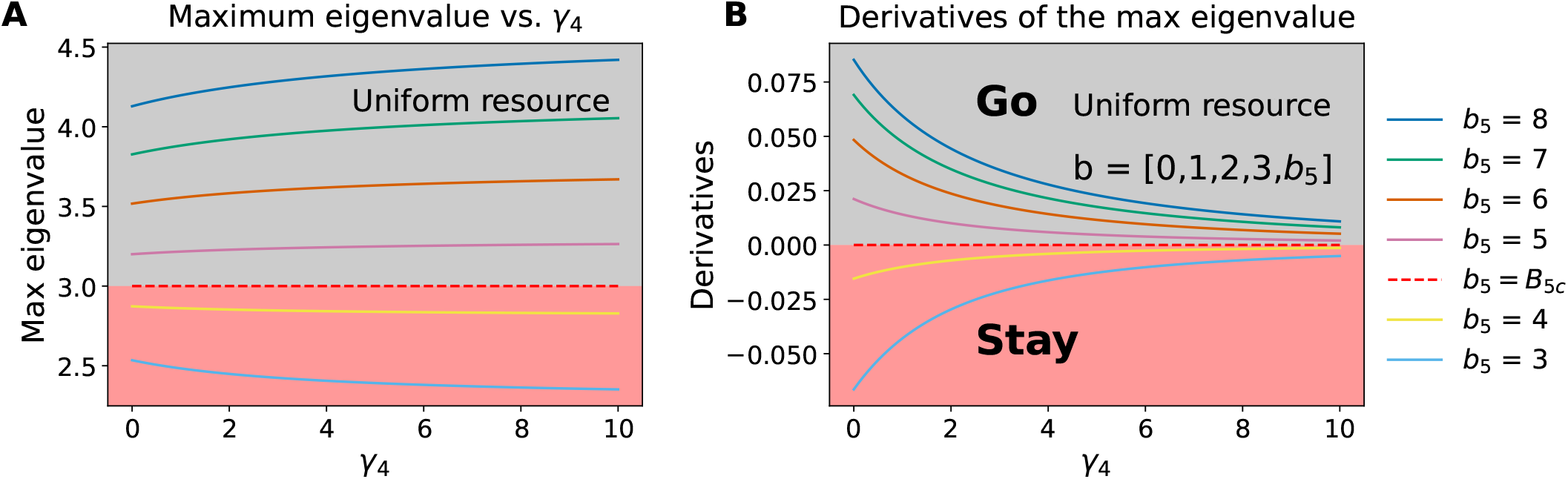
Maximum eigenvalue Λ_*γ*_ as an effective objective function. **A.** The maximum eigenvalue Λ_*γ*_ of the low-density coefficient matrix 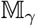 predicts the most competitive foraging strategies *γ* _opt_. Typically, the optimal values are *γ*_1,2,3_ = 10 and *γ*_5_ = 0, but the value of *γ*_4_ depends on the birth rate at resource level *S*_5_, *b*_5_. When *b*_5_ is above the critical value *B*_5*c*_, the phenotype with *γ*_4_ → *γ*_*max*_ has the largest maximum eigenvalue and thus are predicted to be the optimal; however when *b*_5_ is below *B*_5*c*_, the phenotype with *γ*_4_ → *γ*_*min*_ is predicted to dominate the population instead. The same transition in the full agent-based stochastic simulation is shown in Fig. 3. **B.** The derivatives of the eigenvalue predict a range of *γ*_4_ values yield similar maximum eigenvalues when *b*_5_ is close to *B*_5*c*_. The effective objective function is insensitive to changes of *γ*_4_, resulting in a large variability of *γ*_4_ as seen in the simulation results shown in Fig. 2E and F. When the derivative is positive, larger *γ*_4_ leads to larger *λ*_*γ*_ and thus agents are better off leaving their current location to explore (”go”). When the derivative is negative, smaller *γ*_4_ leads to larger *λ*_*γ*_ and thus agents are better off exploiting their current location (”stay”).

### Violin plots show the phenotype coexistence at a range of densities

While it is possible to derive an effective objective function in the low-density limit, increasing agent density invalidates this approximation and precludes the derivation of a well-defined objective function of only a single phenotype vector *γ*. It is observed from the full agent-based stochastic simulation that there exists a range of initial density values with which different phenotypes coexist for an evolutionarily long time, at least up to 20000 iterations of the full stochastic simulations shown in Fig. 4. Here we present the violin plots of the relocation rate distributions of the full agent-based stochastic simulations at different initial densities in Supplementary Fig. 2.

**FIG. 2.**
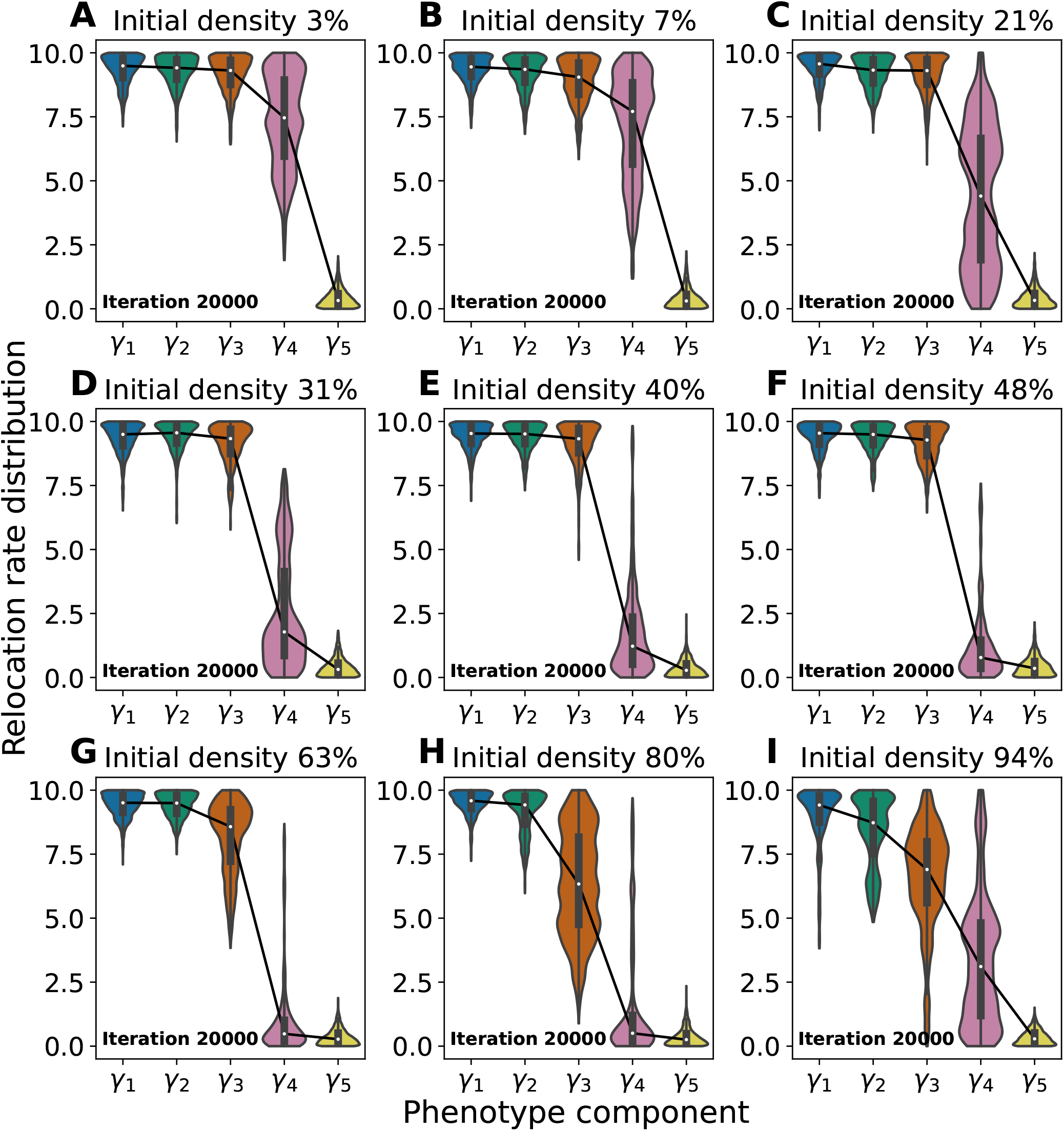
Phenotype distributions of agent-based stochastic simulations at different initial densities. The violin plots of the hopping rate distributions at the end of the agent-based stochastic simulations (i.e., at iteration 20000) show the coexistence of phenotypes at a range of initial densities, indicated by a large width of the distribution. At lower densities, coexistence occurs when *b*_5_ ≈ *B*_5*c*_, while some degree of coexistence is observed for several initial densities without tuning *b*_5_. Note that the large degree of coexistence at 94% density is mostly due to the environment being so dense that opportunities for relocation and birth are extremely rare.

### The well-mixed assumption introduces a mismatch in timescales

The full agent-based model has a population of agents navigating on a 2*d* environment. For analytical tractability, this full model is reduced to a 5-state model that eliminates spatial dependencies. However, it is observed in Fig. 2E and F that the full model and the reduced model have a qualitative mismatch in the order of magnitude of the convergence speed, as well as a quantitative mismatch in the value of the transition point of *b*_5_. Here in Supplementary Fig. 3 we show the comparison of the two models at a shorter timescale (700 iterations) with different *b*_5_ values (4 and 5). The results suggest that the well-mixed assumption introduces the discrepancy in timescales, as simulations of a stochastic 5-state model agree with the mean-field 5-state model.

**FIG. 3.**
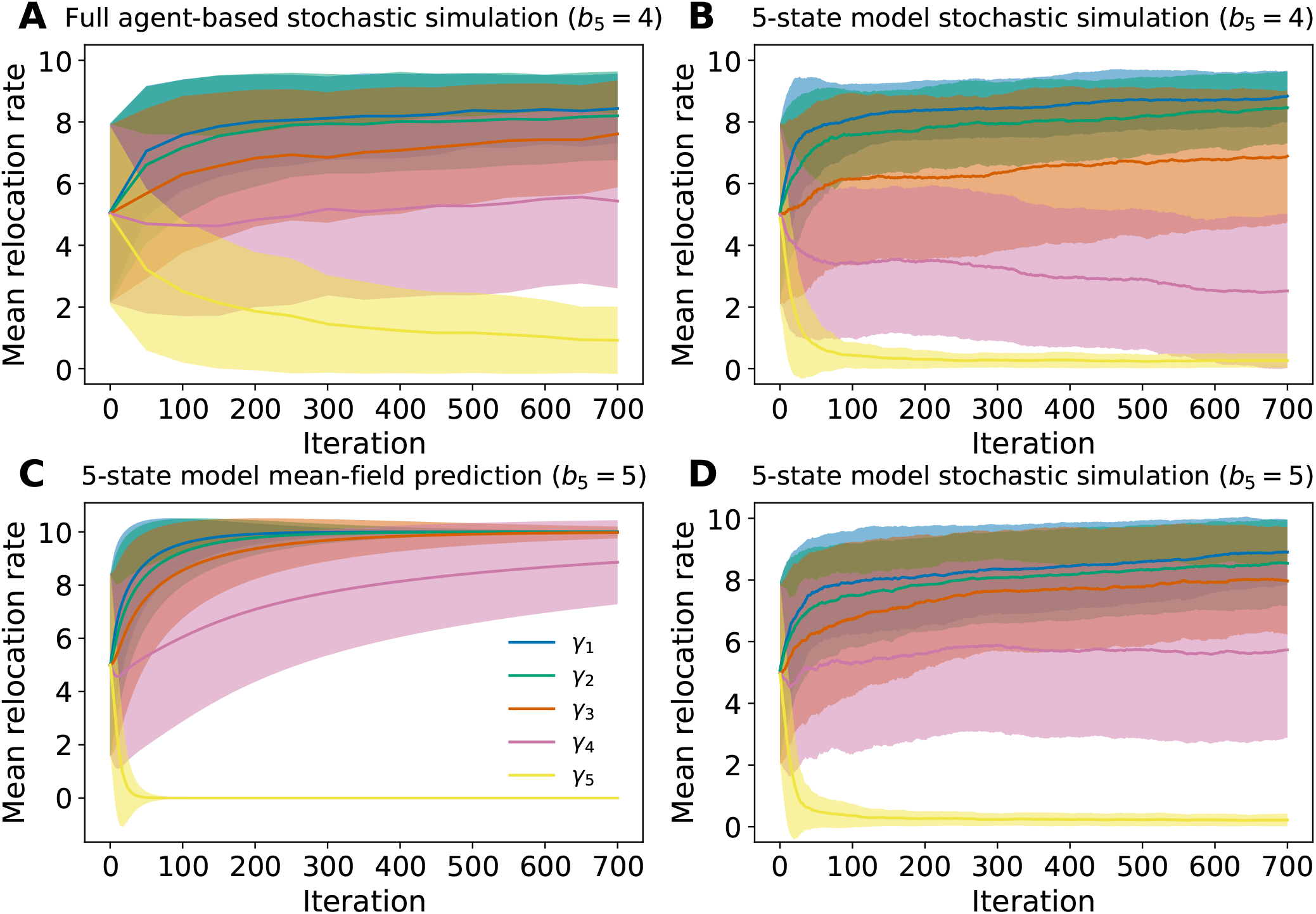
Differences between the full agent-based model and the minimal 5-state model. **A.** The mean relocation rates of the full 2*d* stochastic simulation with *b*_5_ = 4 have not converged by 700 catastrophes. **B.** Agent-based simulations of a 5-state model with *b*_5_ = 4 show a faster convergence of *γ*_5_ to 0 and *γ*_1,2,3_ to high values near *γ*_max_ = 10, within the same 700 catastrophes shown in A (note the different scales of the horizontal axes in Fig. 2E and F). **C.** The 5-state mean field model with *b*_5_ = 5 (which predicts *γ*_4_ → 10 at long times) converges within 700 catastrophes, though the fate of *γ*_4_ is different from the stochastic 5-state model with *b*_5_ = 4. **D.** The stochastic agent-based 5-state model with *b*_5_ = 5 largely converges within 700 iterations and agrees slightly better with the mean-field model’s prediction for *γ*_4_.

## References

[1] D. W. Stephens, J. S. Brown, and R. C. Ydenberg, Foraging: behavior and ecology (University of Chicago Press, 2008).

[2] R. G. P. P. Marra, Birds of two worlds: the ecology and evolution of migration (JHU Press, 2005).

[3] A. López-Cruz, A. Sordillo, N. Pokala, Q. Liu, P. T. McGrath, and C. I. Bargmann, Neuron 102, 407 (2019).

[4] M. Doebeli, Y. Ispolatov, and B. Simon, Elife 6, e23804 (2017).

[5] J. Zylberberg and M. R. DeWeese, Frontiers in computational neuroscience 5, 20 (2011).

[6] Z. P. Kilpatrick, J. D. Davidson, and A. E. Hady, “Normative theory of patch foraging decisions,” (2020), arXiv:2004.10671 [q-bio.NC].

[7] Z. P. Kilpatrick, J. D. Davidson, and A. E. Hady, bioRxiv (2021), 10.1101/2021.04.24.441241, https://www.biorxiv.org/content/early/2021/04/26/2021.04.24.441241.full.pdf.

[8] P. C. Bressloff, Phys. Rev. E 103, 012101 (2021).

[9] I. M. Park and J. W. Pillow, BioRxiv, 178418 (2017).

[10] W. Młynarski, M. Hledík, T. R. Sokolowski, and G. Tkačik, Neuron 109, 1227 (2021).

[11] E. D. Karpas, A. Shklarsh, and E. Schneidman, Proceedings of the National Academy of Sciences 114, 5589 (2017).

[12] S. B. Laughlin, Z Naturforsch 36c, 910 (1981).

[13] W. Bialek and W. G. Owen, Biophysical journal 58, 1227 (1990).

[14] R. d. R. Van Steveninck, W. Bialek, M. Potters, and R. Carlson, in Proceedings of IEEE International Conference on Systems, Man and Cybernetics, Vol. 1 (IEEE, 1994) pp. 302–307.

[15] L. M. Bautista, J. Tinbergen, P. Wiersma, and A. Kacelnik, The American Naturalist 152, 543 (1998).

[16] A. Pérez-Escudero, M. Rivera-Alba, and G. G. de Polavieja, Proceedings of the National Academy of Sciences 106, 20544 (2009).

[17] A. Celani and M. Vergassola, Proceedings of the National Academy of Sciences 107, 1391 (2010), https://www.pnas.org/content/107/4/1391.full.pdf.

[18] J. Diedrichsen, R. Shadmehr, and R. B. Ivry, Trends in Cognitive Sciences 14, 31 (2010).

[19] J. V. Rodrigues and E. I. Shakhnovich, eLife 8, e50509 (2019).

[20] S. Powell, S. L. Price, and D. J. C. Kronauer, Proceedings of the National Academy of Sciences 117, 6608 (2020), https://www.pnas.org/content/117/12/6608.full.pdf.

[21] S. W. Buskirk, A. B. Rokes, and G. I. Lang, eLife 9, e62238 (2020).

[22] A. J. Calhoun, S. H. Chalasani, and T. O. Sharpee, eLife 3, e04220 (2014).

[23] D. T. Gillespie, Annu. Rev. Phys. Chem. 58, 35 (2007).

[24] N. Goldenfeld, Lectures on Phase Transitions and the Renormalization Group (Westview Press, 1992).

[25] M. Assaf and B. Meerson, Journal of Physics A: Mathematical and Theoretical 50, 263001 (2017).

[26] J. Lehman, J. Clune, D. Misevic, C. Adami, L. Altenberg, J. Beaulieu, P. J. Bentley, S. Bernard, G. Beslon, D. M. Bryson, et al., Artificial life 26, 274 (2020).

[27] R. P. Mann, PLOS Computational Biology 17, 1 (2021).

[28] R. S. Sutton and A. G. Barto, Reinforcement learning: An introduction (MIT press, 2018).

[29] I. M. Dobbs and I. Molho, Journal of Evolutionary Economics 9, 187 (1999).

[30] T. L. Martin and R. B. Huey, The American Naturalist 171, E102 (2008), pMID: 18271721, https://doi.org/10.1086/527502.

[31] E. Vercken, M. Wellenreuther, E. I. Svensson, and B. Mauroy, PLOS ONE 7, 1 (2012).

[32] B. Confavreux, F. Zenke, E. Agnes, T. Lillicrap, and T. Vogels, in Advances in Neural Information Processing Systems, Vol. 33, edited by H. Larochelle, M. Ranzato, R. Hadsell, M. F. Balcan, and H. Lin (Curran Associates, Inc., 2020) pp. 16398–16408.

[33] E. Najarro and S. Risi, in Advances in Neural Information Processing Systems, Vol. 33, edited by H. Larochelle, M. Ranzato, R. Hadsell, M. F. Balcan, and H. Lin (Curran Associates, Inc., 2020) pp. 20719–20731.

[34] J. Jordan, M. Schmidt, W. Senn, and M. A. Petrovici, “Evolving to learn: discovering interpretable plasticity rules for spiking networks,” (2021), arXiv:2005.14149 [q-bio.NC].

[35] H. D. Mettler, M. Schmidt, W. Senn, M. A. Petrovici, and J. Jordan, “Evolving neuronal plasticity rules using cartesian genetic programming,” (2021), arXiv:2102.04312 [cs.NE].

[36] F. Mery and J. G. Burns, Evolutionary Ecology 24, 571 (2010).

[37] I. C. G. Kadow, Current Opinion in Neurobiology 54, 60 (2019).

[38] G. E. Robinson and A. B. Barron, Science 356, 26 (2017).

[39] P. C. Bressloff, Journal of Physics A: Mathematical and Theoretical 50, 133001 (2017).

[40] F. Jabot and J. Chave, The American Naturalist 178, E37 (2011).

[41] B. Haegeman and M. Loreau, Journal of theoretical biology 269, 150 (2011).

[42] S. Azaele, S. Suweis, J. Grilli, I. Volkov, J. R. Banavar, and A. Maritan, Rev. Mod. Phys. 88, 035003 (2016).

[43] A. Bazzani, G. Castellani, E. Giampieri, and C. Sala, Master Equation and Relative Species Abundance Distribution for Lotka-Volterra Models of Interacting Ecological Communities, 37 (2016).

[44] T. Butler and D. Reynolds, Phys. Rev. E 79, 032901 (2009).

[45] T. Butler and N. Goldenfeld, Phys. Rev. E 80, 030902 (2009).

[46] H.-Y. Shih and N. Goldenfeld, Phys. Rev. E 90, 050702 (2014).

[47] C. Xue and N. Goldenfeld, Phys. Rev. Lett. 119, 268101 (2017).

[48] U. Dieckmann and R. Law, Journal of mathematical biology 34, 579 (1996).

[49] U. Dobramysl, M. Mobilia, M. Pleimling, and U. C. Täuber, Journal of Physics A: Mathematical and Theoretical 51, 063001 (2018).

[50] J. T. Wootton, Nature 433, 309 (2005).

[51] D. Alonso, R. S. Etienne, and A. J. McKane, Trends in ecology & evolution 21, 451 (2006).

[52] O. Allouche and R. Kadmon, Journal of Theoretical Biology 258, 274 (2009).

[53] E. K. Agarwala, H. J. Chiel, and P. J. Thomas, Journal of theoretical biology 304, 235 (2012).

[54] R. Harpaz and E. Schneidman, Elife 9, e56196 (2020).

[55] S. S. Chow, C. O. Wilke, C. Ofria, R. E. Lenski, and C. Adami, Science 305, 84 (2004).

[56] B. J. McGill and J. S. Brown, Annu. Rev. Ecol. Evol. Syst. 38, 403 (2007).

[57] R. S. Olson, A. Hintze, F. C. Dyer, D. B. Knoester, and C. Adami, Journal of The Royal Society Interface 10, 20130305 (2013).

[58] C. Adami, J. Schossau, and A. Hintze, Physics of life reviews 19, 1 (2016).

[59] D. Goldberg, (1989).

[60] F. I. Thomas, L. T. Kregting, B. D. Badgley, M. J. Donahue, and P. O. Yund, Marine Ecology Progress Series 494, 231 (2013).

[61] H. Murua, Environmental biology of fishes 97, 329 (2014).

[62] S. H. Alonzo, K. A. Stiver, and S. E. Marsh-Rollo, Nature Communications 7, 1 (2016).

[63] Y. H. Hussain, J. S. Guasto, R. K. Zimmer, R. Stocker, and J. A. Riffell, Journal of Experimental Biology 219, 1458 (2016).

[64] K. Spoelstra, M. Wikelski, S. Daan, A. S. I. Loudon, and M. Hau, Proceedings of the National Academy of Sciences 113, 686 (2016), https://www.pnas.org/content/113/3/686.full.pdf.

[65] C. Deroulers and R. Monasson, Physical Review E 69, 016126 (2004).

[66] D. Shaikh and I. Rañó, Frontiers in Bioengineering and Biotechnology 8(2020).

[67] S. Dvoretskii, Z. Gong, A. Gupta, J. Parent, and B. Alicea, arXiv preprint arXiv:2003.07689 (2020).

[68] C. J. Cueva and X.-X. Wei, in International Conference on Learning Representations (2018).

[69] C. R. Harris, K. J. Millman, S. J. van der Walt, R. Gommers, P. Virtanen, D. Cournapeau, E. Wieser, J. Taylor, S. Berg, N. J. Smith, R. Kern, M. Picus, S. Hoyer, M. H. van Kerkwijk, M. Brett, A. Haldane, J. Fernández del Río, M. Wiebe, P. Peterson, P. Gérard-Marchant, K. Sheppard, T. Reddy, W. Weckesser, H. Abbasi, C. Gohlke, and T. E. Oliphant, Nature 585, 357–362 (2020).

[70] P. Virtanen, R. Gommers, T. E. Oliphant, M. Haberland, T. Reddy, D. Cournapeau, E. Burovski, P. Peterson, W. Weckesser, J. Bright, S. J. van der Walt, M. Brett, J. Wilson, K. J. Millman, N. Mayorov, A. R. J. Nelson, E. Jones, R. Kern, E. Larson, C. J. Carey, İ. Polat, Y. Feng, E. W. Moore, J. VanderPlas, D. Laxalde, J. Perktold, R. Cimrman, I. Henriksen, E. A. Quintero, C. R. Harris, A. M. Archibald, A. H. Ribeiro, F. Pedregosa, P. van Mulbregt, and SciPy 1.0 Contributors, Nature Methods 17, 261 (2020).

[71] J. D. Hunter, IEEE Annals of the History of Computing 9, 90 (2007).

[72] E. Bonabeau, Proceedings of the national academy of sciences 99, 7280 (2002).

[73] R. A. Horn and C. R. Johnson, Matrix Analysis, 2nd ed. (Cambridge University Press, USA, 2012).

